# Rapid neocortical network modifications via dendritic plateau potential induced plasticity

**DOI:** 10.1101/2025.11.19.689338

**Authors:** Kuo Xiao, Yiding Li, Brennan J. Sullivan, Guanchun Li, Jeffrey C. Magee

**Affiliations:** Howard Hughes Medical Institute, Baylor College of Medicine, Houston, TX; Jan and Dan Duncan Neurological Research Institute, Texas Children’s Hospital, Houston, TX

**Author notes:** Correspondence to: Kuo Xiao, HHMI/Baylor College of Medicine, Jan and Dan Duncan Neurological Research Institute, 1250 Moursund Dr., Houston, TX, 77030, USA. Equal contribution (Kuo Xiao, Yiding Li, Brennan J. Sullivan, Guanchun Li).

## Abstract

Learning in brains is associated with changes in neuronal network activity thought to be driven by synaptic plasticity. While recent work in the hippocampus has revealed some of the mechanisms involved there, less is known about how neocortical circuits adapt, especially during behavior. Here to determine if neocortical areas possess rapid plasticity mechanisms that could support online adaptations we used optical imaging and intracellular membrane potential (Vm) recordings to examine the activity of layer V neurons in a higher visual area of mice learning a task. The introduction of a novel rewarded stimulus resulted in a rapid modification of population activity that featured abrupt alterations in single neuron selectivity. Vm recordings revealed that both naturally occurring and experimentally-induced dendritic calcium plateau potentials (plateaus) rapidly alter the action potential (APs) output and Vm dynamics of neurons over many seconds of time around the plateau, in some cases from one trial to the next. Trains of high frequency APs had no effect. Finally, experimental inhibition during learning of the distal dendritic region responsible for initiating plateaus reduced the rate of population level adaptation. Our findings suggest that the deep layers of higher order visual cortex possess a rapid learning mechanism mediated by plateau-induced synaptic plasticity.

## Main

Brains adapt their neuronal representations and dynamics to support learned behaviors and cognitive functions. This process is mediated by learning-related changes in neuronal network population activity and alterations in the strength of synaptic connections between neurons in these networks may play an important role. Indeed, a synaptic plasticity form known as Behavioral Timescale Synaptic Plasticity (BTSP) has been recently reported to underlie the adaptive changes in the hippocampus that are required for learning from experience^1–8^. There, BTSP is driven by long-duration (>100 ms) dendritic calcium plateau potentials (plateau) that adjust the weights of active synaptic inputs over many seconds of time to generate new place cell activity. The unique properties of plateau driven synaptic plasticity, which include rapid, one-shot changes linked to both current experience and past learning^9,10^, suggest a potentially powerful learning and memory mechanism that could be present in many brain regions.

Neocortical networks are composed of a pyramidal neuron-based circuit motif that is similar to that present in the hippocampus^1,11^. Evidence suggests that neuronal activity in various neocortical areas is a complex mixture of multiple features that enable context-dependent population level representations^12–18^. There are reports that this population activity can undergo substantial modification during specific learning tasks^19–26^ and *in vitro* work has revealed the presence of synaptic plasticity mechanisms (mostly Hebbian forms) within different neocortical areas^1,27,28^. However, there are also both theoretical and experimental studies that hold the neocortex to be a slow learning system that mostly changes during offline periods of re-activation, primarily during sleep^29–32^. These studies are based on the ideas that there should be less need for rapid plasticity in the neocortex since the fundamental characteristics of the external, and perhaps internal, worlds do not substantially change over short time frames and that brains should possess a large capacity storage area that is less plastic for the long-term maintenance of important information (i.e memory). Thus, we sought to explore whether neocortical regions possess rapid plasticity mechanisms that could allow online adaptations of population activity.

### Rapid network activity changes

We began by recording a population of several hundred layer V (L5) neurons that we estimate to be in the lateral visual areas (primarily laterointermediate (LI) or lateromedial (LM)) of a group of *Rbp4-cre::Ai93* mice (n = 478 cells from 7 mice) expressing GCaMP6f (Fig. 1a, Extended Data Fig. 1a and Methods). We employed a visually-guided spatial navigation task where head-fixed mice were trained to traverse a linear track to a location where a specific visual stimulus (horizontally moving large disc; stimulus 1) indicated the presence of a water reward delivered upon stimulus offset (Fig. 1b,c). Following initial training, we implemented a contingency shift during the experimental recording sessions. After establishing a baseline period with only the familiar stimulus present (block A, trials 1–50), a novel visual stimulus (vertically moving large disc; stimulus 2) was introduced at a different location along the track, and the reward delivery was re-associated exclusively with this new stimulus (block B; trials 51–100). This contingency shift produced a clear behavioral adaptation where mice adjusted their licking behavior such that licking decreased at the old now non-rewarded site and increased at the new stimulus-reward site (Fig. 1c; upper traces). Concurrently, the mice progressively modified their running profile to match the new, reward-paired stimulus location (Fig. 1c; lower traces). Together, these behavioral results suggest that mice actively learned the updated stimulus-reward association when the contingency was shifted, adapting their navigational and reward-seeking strategies accordingly.

**Fig. 1.**
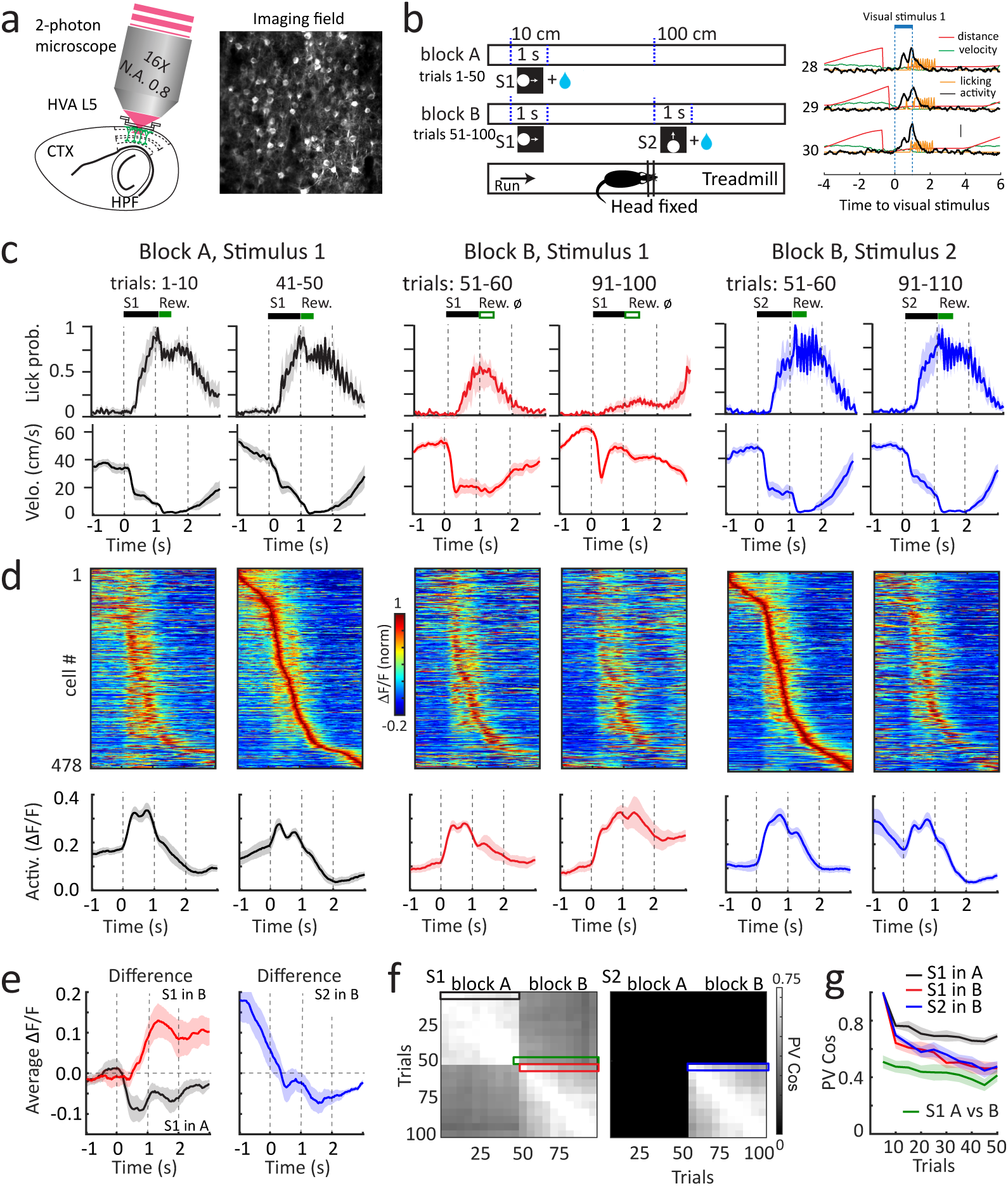
Rapid formation of context-dependent visual representations in HVA. **a,** Experimental setup for two-photon imaging and an example field of view (FOV). **b**, Left, schematic of the behavioral paradigm. Right, three consecutive trails aligned to S1, showing distance (red), velocity (green), licking (orange), and an example cell activity (black). Blue dashed lines indicate the period of S1. Scale bar: 1 m (distance), 1 m/s (velocity), 1 ΔF/F (activity). **c,** Behavioral measurements (licking and velocity) during the first and last 10 trials of each block, aligned to stimuli presentations (solid line, mean; shaded areas, mean ± SEM, n = 7 animals). **d,** Heatmap illustrating mean activities across all neurons during the behavioral phases shown in **c**, sorted by peak activity timing during the last 10 trials of block A (aligned to S1) or the first 10 trials of block B (aligned to S2). Average neuronal activities (solid line, mean; shaded area, mean ± SEM, n = 7 animals) are shown below. **e,** Differences in average activity between first and last 10 trials for each block, aligned to S1 (left; black: block A, red: block B) and S2 (right; blue: block B). **f,** Cosine Similarity of population vectors (PV) across trails, aligned to S1 (left) and S2 (right). **g,** PV cosine similarity (solid line, mean; shaded area, mean ± SEM, n = 7 animals) comparing subsequent trials with the first 5 trials within block A (green, aligned to S1), within block B (red, aligned to S1; blue, aligned to S2), and trials in block B compared to the last 5 trials of block A (black, aligned to S1). Color coding matches the boxes in **f**.

We examined the somatic activity of L5 neurons during the above behavior. Heat maps of mean activity from individual neurons (Fig. 1d; upper) and population averages (Fig. 1d; lower) of neuronal activity indicate that L5 neurons produce a sequence of activity that traverses the entire running lap including regions both inside and outside the visual stimuli (Extended Data Fig. 1b–e). During block A (familiar visual stimulus, S1), the average amplitude of population activity as well as the trial-by-trial population vector correlations (PV cosine similarity) modestly decreased as trials proceeded (Fig. 1e–g; black). However, we observed a large and rapid change in the population level activity upon transition from block A to block B (PV cosine similarity, Fig. 1f,g; green), that progressively continued throughout block B (Fig. 1f,g; red and blue). This population level change associated with the introduction of a novel rewarded stimulus (S2) in block B appeared to be caused by large increases in neuronal activity in the extra-stimulus regions between the two stimuli (Fig. 1e; red 1∼2s and blue −1∼0s) and these changes were correlated with behavioral indices (Extended Data Fig. 1f). In sum, the introduction of a novel rewarded stimulus resulted in a rapid modification of the population activity of L5 neurons in higher visual area (HVA) that featured a prominent increase in neuronal activity in the inter-stimulus period.

To explore the impact of the new rewarded stimulus on the activity of individual L5 neurons we performed a Tensor Component Analysis (TCA) on the neuronal activity organized into a three dimensional array of neurons × trial time × trials across sessions (Extended Data Fig. 2a and Methods; all mice were combined into a single array; 478 neurons from 7 mice)^33^. This unsupervised tensor decomposition of neuronal activity identified seven different components with different groups of neurons whose activity was best described by one of these components (Fig. 2a and Extended Data Fig. 2b,c). Each of these neuronal groups showed unique temporal dynamics relative to visual stimulus presentation, changes occurring across trials, and the proportions of neurons most strongly associated with each TCA component (Fig. 2a). Notably, only the two components associated with activity in the period between S1 and S2 (Fig. 2a–c, classes 4 & 5) showed increases in the weight amplitude of the associated components across trials during block B. These data suggest that the activity of the neurons in these two groups increased throughout block B and this was confirmed by examining the actual activity of all neuronal groups separately (Fig. 2b,c and Extended data Fig. 2d). Furthermore, in many cases the activity of individual neurons changed abruptly and in some cells from one trial to the next following the transition from block A to block B (see examples in Fig. 2d,e and Extended data Fig. 2e).

**Fig. 2.**
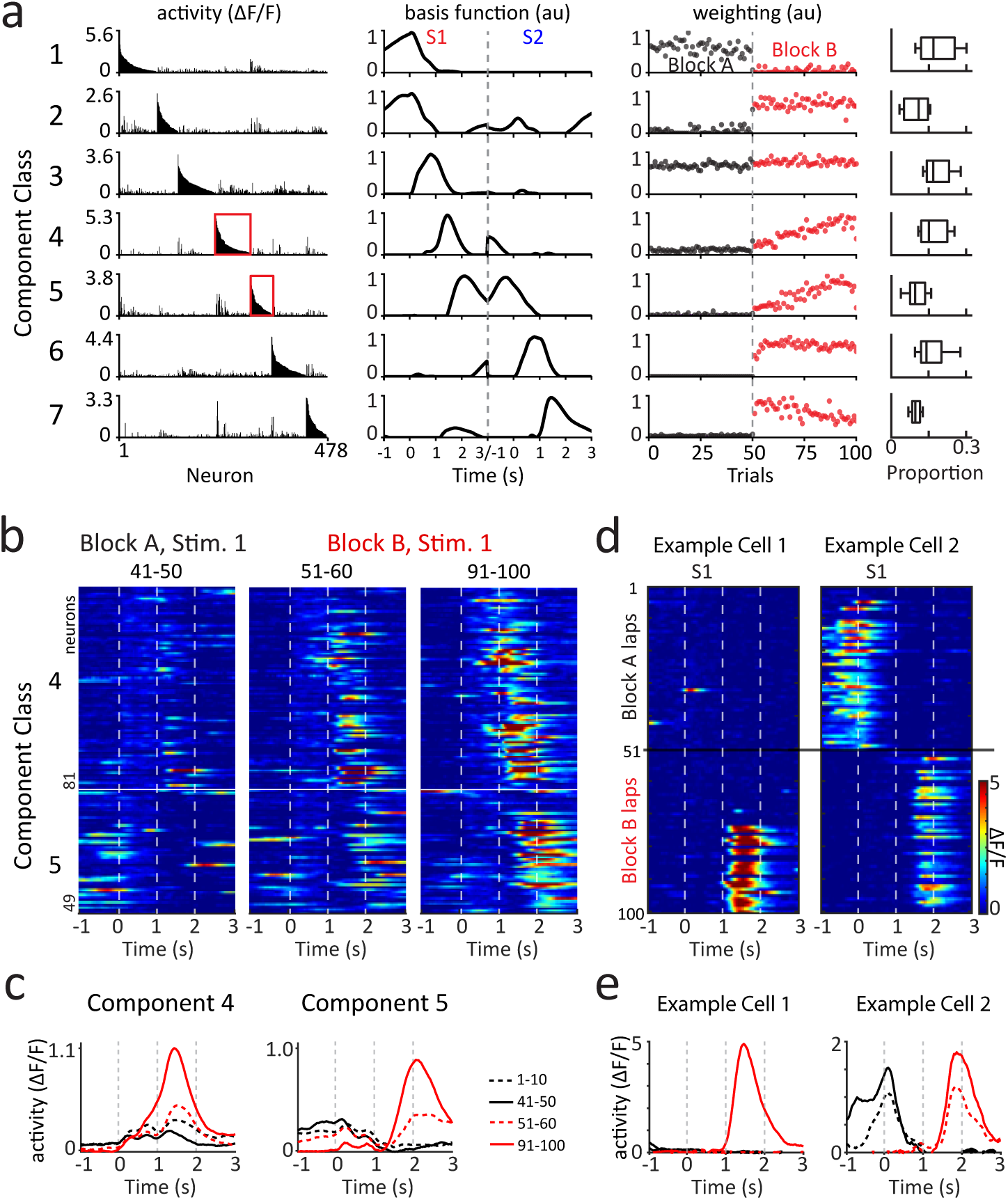
Abrupt changes in single neuron feature selectivity. **a,** Seven low-dimensional components identified by a rank-7 Tensor Component Analysis (TCA) model. Each component contains a neuron factor (left column), temporal basis function (middle column), and trial weighting factor (right column). The trial weighting factor is color-coded according to the block assignment of each trial. Different components capture unique neural dynamics both within different periods relative to visual stimulus presentation and across trials. Boxplots to the right illustrate the proportions of neurons most strongly associated with each TCA component (n = 7 animals). **b,** Mean activities of all neurons from components 4 and 5 in **a** during the first 10 or last 10 trials of each block, aligned to S1 and sorted according to cell’s component association from the TCA model shown in **a**. **c**, Average activities for neurons associated with component 4 and 5 in **b**. **d,** Activity of representative neurons associated with components 4 and 5 across all trials in both blocks. **e,** Average activities of representative neurons in **d** during different trials in both blocks.

To further quantify the above changes in neuronal activity, we constructed a Generalized Linear Model (GLM) to predict neural activity based on a set of behavioral and visual stimulus variables (Extended Data Fig. 3 and Methods; variables including distance run from reward, velocity, licking, extra-stimulus reward, extra-stimulus reward omission, S1 and S2). Fitting the GLM to individual neuron activity produced in many neurons a combination of unique response profiles for different behavior and stimulus variables in both first and last 10 trials of block B (Extended Data Fig. 3d), through which we computed the response amplitude as a quantification of each neuron’s feature selectivity for the variables across different trials (Extended Data Fig. 3e). We found that the neurons associated with the two TCA components active between S1 and S2 showed a nearly three-fold increase in response amplitude (Extended Data Fig. 3f). Thus, individual neurons showed preferred feature selectivity for one of the two visual stimuli or extra-stimulus regions and this selectivity, especially for the extra-stimulus regions, was frequently observed to change following the presentation of a novel rewarded stimulus. We conclude from the above data that there are rapid learning-related modifications in the feature selectivity of individual L5 neurons within this HVA that produce significant alterations in the population activity of the network leading to context-dependent visual representations.

### Plateau potentials and rapid plasticity

That the above neuronal activity modifications occurred over such a short amount of time (∼10 mins), with activity changing even from one trial to the next in some neurons, suggests that a form of rapid plasticity may be present in HVA L5 neurons. Because such rapid plasticity has been observed elsewhere to be induced by plateaus, we next turned to intracellular membrane potential (Vm) recordings from deep-layer HVA putative pyramidal neurons to determine if such electrical signals are present during this behavior (Methods, Fig. 3a and Extended Data Fig. 4a). To begin we characterized the action potential (AP) firing rates and subthreshold Vm dynamics of a set of neurons during task performance (n = 94 neurons; Fig. 3b,c and Extended Data Fig. 4b, 5). We observed that a large majority of the recorded neurons (∼90%) exhibited clear AP and Vm responses to the visual stimuli (Fig. 3b,c and Extended Data Fig. 5) with most showing increases in firing and depolarizing responses (∼70%) and others that exhibited decreases in AP rate and a hyperpolarizing response (∼30%). As observed in the above imaging data, a sequence of AP activity and peak Vm polarization was found across the entire running lap that spanned the time around the familiar stimulus as well as periods clearly outside of the stimulus (−1∼3s stimulus onset; Extended Data Fig. 5c,d).

**Fig. 3.**
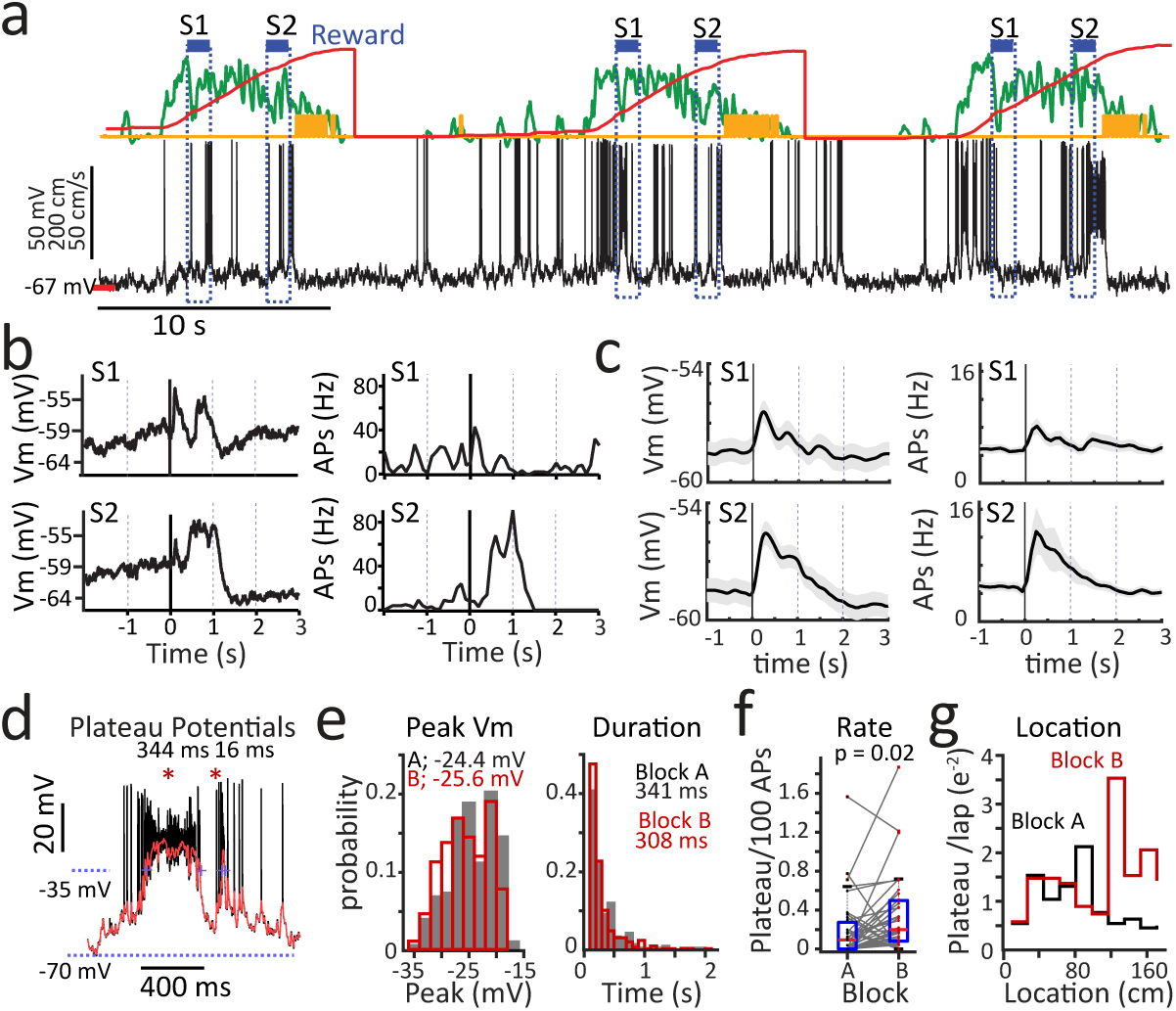
Context changes increase dendritic calcium plateau potential initiation. **a,** Representative traces showing Vm (black), mouse location (red), running speed (green), and licking (yellow ticks). **b,** Average Vm (10 trials) around S1 (top left), S2 (bottom left) and AP rate around S1 (top right), S2 (bottom right) during block B of example cell. **c**, Population average responses for all recorded cells (n = 94 cells) during block B before induction. Panels show average Vm aligned to S1 (top left) and S2 (bottom left), and average AP firing rate aligned to S1 (top right) and S2 (bottom right). **d,** Example plateau potentials (raw Vm; black), smoothed Vm (red). Detection threshold indicated by dash blue line. **e,** distribution of detected plateau amplitude (left) or duration (right) during block A (gray) and block B (red). **f,** plateau rate for all cells with detected plateaus in block A and block B. *p* = 0.0167, n = 35. **g,** Plateau rate versus location for block A (black) and block B (red).

Spontaneous plateaus, including those with the large amplitude and long duration required to produce Vm plasticity in the hippocampus, were observed in a large fraction of the recorded neurons (∼50%; 50 of 94 neurons; Fig. 3d,e). We quantified the rate of plateau occurrence across the different behavioral contexts and found that upon switching the stimulus-reward contingency to associate the reward with the novel stimulus (block B), the overall rate of plateau generation increased significantly compared to block A (0.20 ± 0.055 vs 0.36 ± 0.069; Fig. 3f). This was particularly obvious in a subset of neurons that displayed highly reliable plateau generation specifically linked to the novel stimulus on nearly every trial immediately following the contingency shift (Extended Data Fig. 6a). The occurrence of plateaus in the above neurons exhibited clear spatial modulation along the track. During the initial context (block A) plateaus were predominantly observed near the location of the familiar stimulus and the associated reward site but following the contingency shift (block B), plateau generation became concentrated around the location of the novel stimulus and as well as during the period between the stimuli (Fig. 3g). In contrast, in a separate set of recordings from mice that were stationary throughout the experiment, the fraction of neurons observed with spontaneous plateau was low (∼7%; 3 of 41 neurons; Extended Data Fig. 6b–j) further suggesting behavioral state may play a role in plateau initiation.

We next attempted to determine if the spontaneous initiation of long duration plateaus (>100 ms) was associated with changes in Vm dynamics and AP firing rates. In those neurons where spontaneous long-duration plateaus occurred at a low enough frequency and consistent enough location to allow measurement of changes we observed that single plateaus significantly altered the neuronal AP activity and Vm profile (n = 14 plateaus in 12 neurons; Fig. 4a–d and Extended Fig. 7). On average the occurrence of a single plateau was associated with a significant increase in the AP firing rate (Fig. 4c; upper) and Vm depolarization (Fig. 4c; lower) for several seconds around the plateaus in subsequent trials compared with those trials before the plateaus (Extended Data Fig. 7b,c). Outside of this potentiation region Vm hyperpolarization and decreases in AP firing were observed for an additional several seconds in each direction. In addition, the amplitude and sign of the Vm changes were significantly correlated with the level of Vm polarization present before the plateaus (Fig. 4d). These data suggest that single plateaus induce a bidirectional synaptic plasticity that depends on the initial weight of the synapses.

**Fig. 4.**
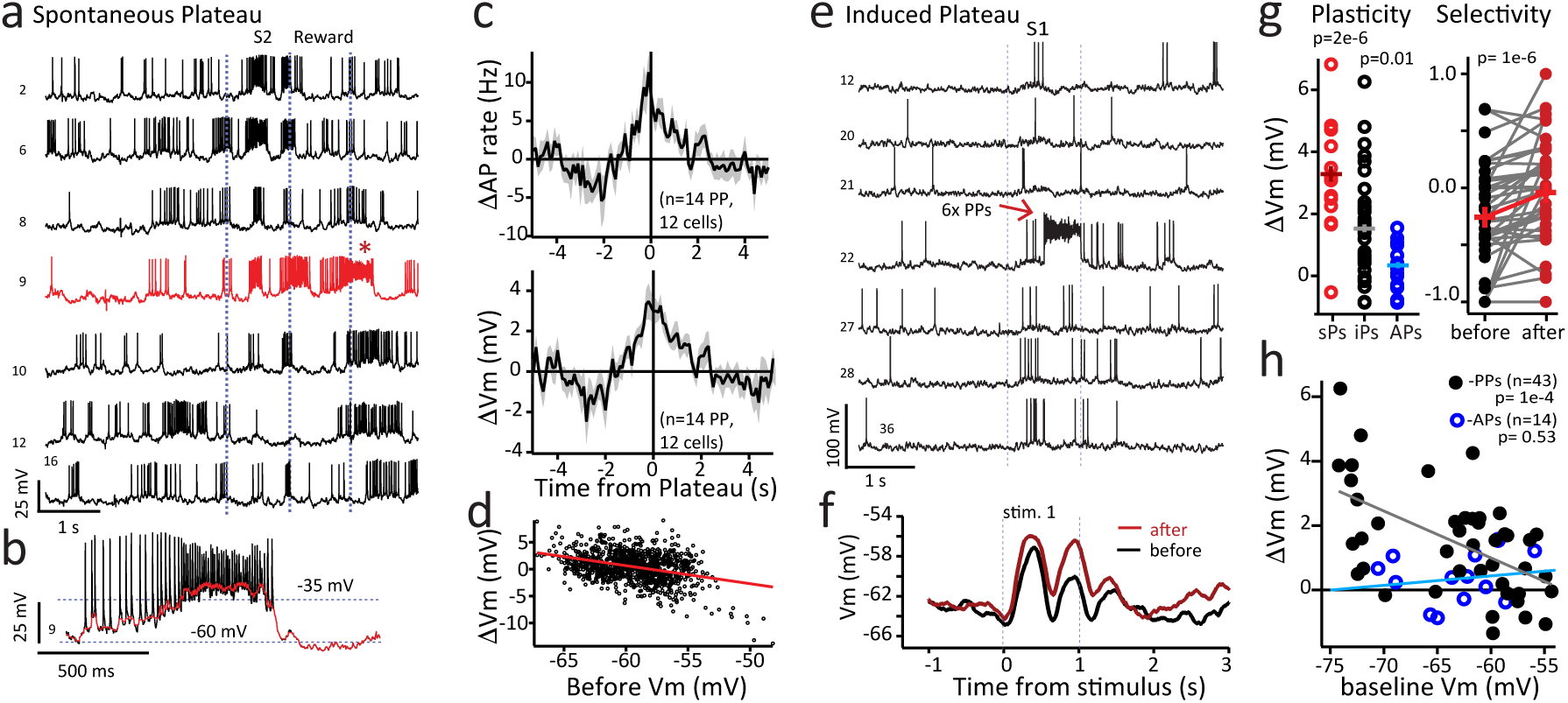
Natural and experimentally induced dendritic calcium plateau potential initiation rapidly alters Vm. **a,** Representative Vm traces for selected laps before (laps 2**–**8) during (red trace, lap 9) and after (laps 10**–**16) first plateau. **b,** Expanded Vm traces of plateau from above trial. **c,** Changed AP rate (upper) and Vm (lower) for trials before and after first plateau. **d,** Average change in Vm versus baseline for population (traces in 100 ms bins, ±4.5 s around plateau, n = 1186 total bins). **e,** Representative Vm traces for selected laps before (laps 12**–**21) during (1 of 6 laps w/ plateaus) and after (laps 27**–**36) induced plateaus. **f,** Average Vm around S1 for 10 trials before (black) and 10 after (red) induced plateaus from cell in **e**. **g,** Right: the change of membrane potential before and after the spontaneous plateaus (sPs), induced plateaus (iPs) and AP induction control (APs). Left: Selectivity index before (black) and after (red) induced PPs (left). *p*-value from two-way, paired students t-test (n = 39). **h,** Change in Vm during plateau induction (black circles; n = 43) and AP induction control (blue circles; n = 14) versus baseline Vm for all neurons. *p*-values from linear correlation.

Motivated by observations suggesting spontaneous plateau could be associated with changes in Vm, we next directly tested the capacity of plateaus to induce Vm plasticity in deep-layer HVA neurons using a controlled protocol. We evoked plateaus in up to six consecutive trials via somatic current injection (∼1 nA, 500 ms), temporally pairing these induced events with S1 (Fig. 4e and Extended Data Fig. 8). Neurons subjected to the plateau induction protocol exhibited a significant potentiation of their Vm response specifically to S1 that was paired with the induction (Fig. 4f). Indeed, this potentiation was input-specific in that the Vm response to the unpaired S2 did not significantly change following plateau induction (Extended Data Fig. 8c,d).

Consequently, the selective strengthening of the response to S1 resulted in a significant increase in neuronal AP firing selectivity between S1 and S2 after plasticity induction (Fig. 4g; right). As a control, a separate group of neurons received high-frequency somatic current injections designed to elicit trains of AP without sustained plateaus during S1 (5 trains of ∼1 nA, 3 ms, 50 Hz injections for 500 ms; Extended Data Fig. 9). Cells undergoing the high-frequency stimulation protocol showed no significant change in their Vm response to S1 (Fig. 4g left and Extended Data Fig. 9c). Finally, we examined factors influencing the magnitude of Vm change associated with the induced plateaus and found a significant correlation between the change in Vm response to the S1 and the pre-induction baseline Vm response (Fig. 4h). This suggests that neurons with initially weaker or more hyperpolarized responses to S1 underwent greater potentiation following plateau induction. Notably, this correlation was absent in the high-frequency stimulation control group (Fig. 4h and Extended Data Fig. 9f), further supporting the specificity of the plateau-mediated plasticity effect. Together these data support the above conclusion that plateaus induce a bidirectional synaptic plasticity that depends on the initial weight of the synapses.

### Experimental inhibition of distal dendrites

To further test the hypothesis that plateaus are required for the observed representational changes (Figs. 1,2), we performed an optogenetic inhibition experiment (Fig. 5). We attempted to suppressed plateau generation in the distal tuft dendrites of L5 neurons by applying widefield laser illumination to the cortical surface (layer 1, L1) in a group of *Rbp4-cre::Ai93* mice (n = 457 cells from 7 mice) whose L5 neurons expressed both GCaMP6f and ArchT (Fig, 5a,b and Extended Data Fig. 10a, cre-dependent ArchT expression produced by local virus injection, Methods). This spatially restricted manipulation was designed to inhibit plateau initiation while minimally affecting baseline neuronal activity^14,16,34^(signal amplitude in ArchT positive neurons was reduced ∼20%; Extended Data Fig. 10b–e). The optogenetic inhibition was applied specifically during the inter-stimulus interval (between S1 and S2) for the first 30 trials of block B, followed by 20 trials without manipulation (Fig. 5c). Consistent with previous findings in other cortical regions showing that inhibition of L1 activity impairs task-related behavior^14,35,36^, this manipulation resulted in a significant delay in behavioral adaptation. Control animals rapidly learned to move through the now non-rewarded S1, whereas mice in the manipulation group continued to slow down and lick after S1 (Fig. 5d and Extended Data Fig.11c). This behavioral impairment was accompanied by a delayed development of the new population-level representation associated with the contingency shift. Specifically, the significant increase in population activity during the post-S1 reward omission period—a hallmark of remapping in control animals (Fig. 1d,e)—was absent in the manipulation group (Fig. 5d,e,h and Extended Data Fig. 11). In contrast, neural responses to the rewarded S2 did not differ between groups, indicating that the manipulation did not cause a general suppression of visually evoked activity (Fig. 5d,e). To further dissect these representational differences, we performed the TCA on these data. This analysis revealed that while visually responsive components were comparable between groups, several non-visual components were significantly altered in the manipulation group, exhibiting changes in both the number of contributing neurons and the temporal profiles of their average activity (Fig. 5f,g and Extended Data Fig. 11a,b components 4, 5). Taken together, these results demonstrate that targeted inhibition of L1 dendritic activity during the inter-stimulus period delays both behavioral adaptation and the corresponding modification of non-visual L5 population activity. These findings indicate that plasticity mechanisms mediated by plateaus, perhaps analogous to hippocampal BTSP, play a critical role in the online development of new neocortical representations.

**Fig. 5.**
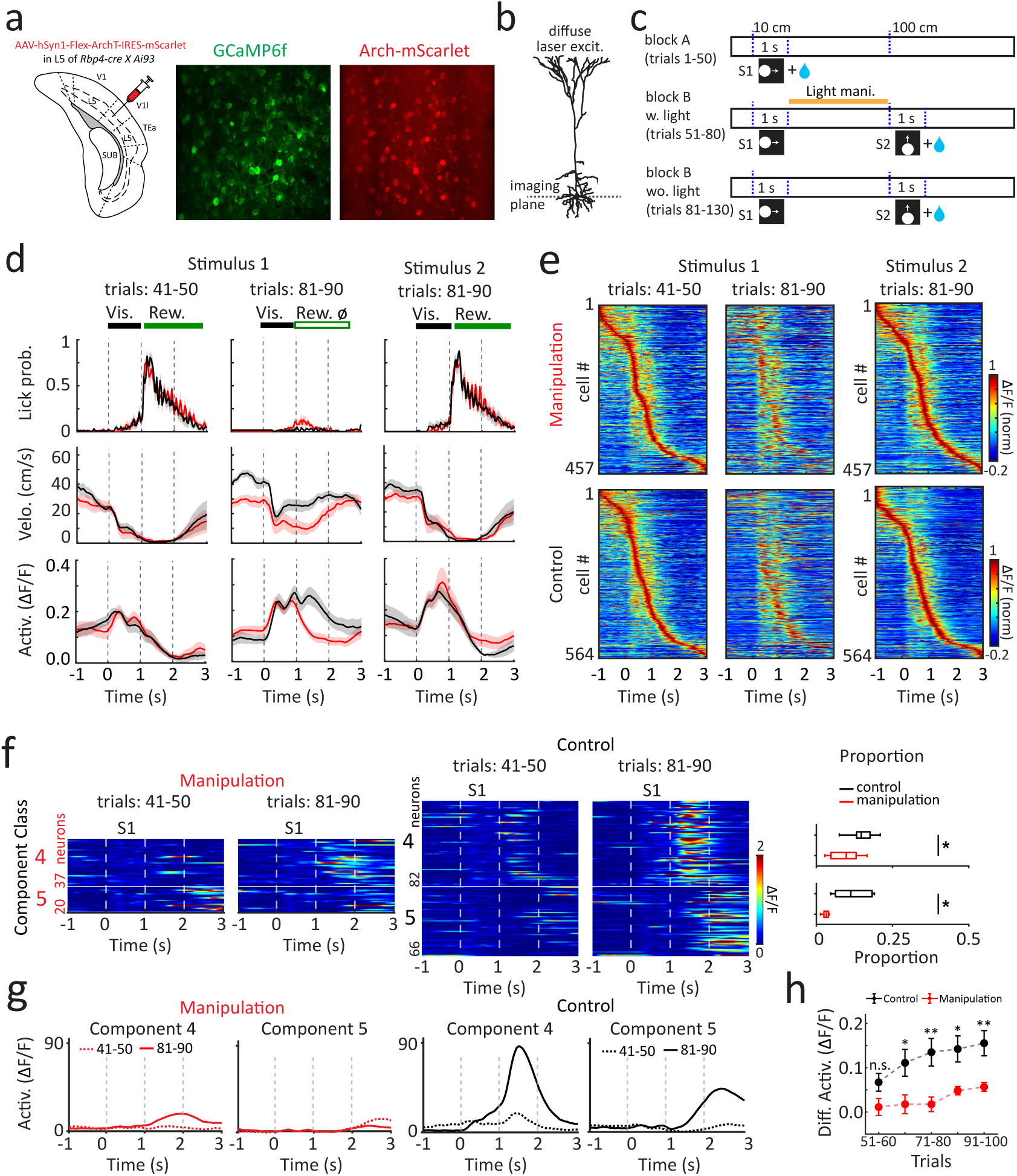
Optogenetic inhibition disrupts rapid neuronal activity modifications. **a**, Illustration of viral injection into L5 of HVA and an example FOV showing Arch-mScarlet expression. **b,** Schematic of optogenetic inhibition. **c,** Schematic of the behavioral paradigm for optogenetic manipulation. The laser was delivered between the two visual stimuli during the first 30 trials of block B. **d,** Behavioral measurements (top: licking; middle: running velocity) and average neuronal activities (bottom) during the last 10 trials of block A and the first 10 trials of block B following optogenetic inhibition, aligned to stimuli onset (black, control group; red, manipulation group; solid line, mean; shaded areas, mean ± SEM, n = 7 animals per group). **e,** Heatmap illustrating mean activities across all neurons during the behavioral phases shown in **d**, sorted by peak activity time during the last 10 trials of block A (aligned to S1) or the first 10 trials of block B following optogenetic inhibition (aligned to S2). **f,** Mean activities of all neurons from TCA components 4 and 5 aligned to S1 for the manipulation and control groups, respectively, during the last 10 trials of block A and the first 10 trials of block B following optogenetic inhibition. Boxplots (right) illustrate the proportions of neurons most strongly associated with each TCA component (black, control group; red, manipulation group; n = 7 animals per group). **g**, Summed neuronal activities of all neurons from component 4 and 5 in **f** across different trials in the manipulation and control groups. **h**, Differences in average activity across all neurons during the 1**–**3 s interval after S1 onset, comparing each behavioral phase to the last 10 trials of block A (black, control; red, manipulation; dot, mean; bar, ± SEM; two-tailed non-paired t-test, *p* = 0.0675, 0.0266, 5.89×10^−3^, 0.0104, 6.70×10^−3^, respectively; n = 7 animals per group).

## Discussion

Here we report that the introduction of a novel rewarded visual stimulus during a visually guided spatial navigation task leads to a rapid and significant modification of the population activity of L5 pyramidal neurons in a HVA that result in context-dependent representations. These changes are associated with, in some cases, abrupt changes in the feature selectivity of individual neurons. In addition, we find that the context changes significantly elevate the occurrence of dendritic calcium plateau potentials and that such plateaus rapidly alter the Vm profile and associated AP firing of individual neurons in a manner consistent with the induction of the synaptic plasticity form known as BTSP in the hippocampus. Additional results from experimentally induced plateau initiation and optogenetic inhibition of the dendritic region where long-duration plateaus are generated support this conclusion. Altogether we present evidence that initiation of single plateau rapidly induces a bi-directional synaptic plasticity that spans many seconds to significantly alter the AP firing selectivity of individual neurons. Furthermore, such plasticity appears to play an important role in the online development of context-dependent representations in a HVA that are useful for spatial navigation.

Our findings are broadly consistent with previous studies demonstrating plasticity within visual neocortical areas during associative learning tasks^19–22,24–26,37–42^. However, most prior work, often focusing on superficial layers (Layer 2/3), primarily described representational changes developing over extended periods, ranging from one week to one month, and focused less on within-session dynamics^19–25^; ^but see 26^. In contrast, our analysis highlights significant representational changes occurring rapidly, within the timescale of a single behavioral block (<10 minutes). That deep-layer HVA populations can undergo rapid adjustments to accommodate new task demands suggests that rapid plasticity mechanisms exist within this neocortical area.

While we have observed such a rapid plasticity mechanism that is induced by plateaus in a HVA, there were notable differences in the properties of this neocortical version and those described previously for BTSP in the hippocampus^2,6–8^. In particular, the magnitude of potentiation induced by our protocol appears more modest in HVA (maximum potentiation; <5 mV in HVA versus <10 mV in CA1). The most obvious explanation would be that some of the input pathways to L5 neurons are less plastic. Along these lines there have been reports that lower NMDA/AMPA ratios in feedforward pathways in visual cortex reduce the level of plasticity expressed at these synapses when compared to other inputs such as the local recurrent inputs^43^. Other issues such as the unknown activity patterns of various input streams to L5 neurons in this region inhibited our ability to determine the exact time-course of plateau-induced plasticity. Thus, additional experiments particularly in brain slices are needed to resolve the exact time course of the underlying synaptic plasticity kernel. Future experiments should also explore which excitatory input pathways are sensitive to plateau-induced plasticity and the roles of other circuit elements such as local inhibitory interneurons and long-range L1 instructive inputs. In addition, the role of neuromodulatory inputs (e.g., cholinergic, dopaminergic states) in permitting or promoting the initiation of dendritic plateau potentials should receive further investigation.

In conclusion, our findings suggest that a plateau-induced plasticity mechanism similar to hippocampal BTSP is present within the deep layers of a HVA. Given the observed properties of this plasticity mechanism it may substantially contribute to the rapid learning-related changes in neuronal responses that we also observed. Altogether the above results support the idea that synaptic plasticity driven by dendritic calcium plateau potentials may function as a general learning mechanism in many brains regions.

## Supplemental Figures

**Extended Data Fig. 1.**
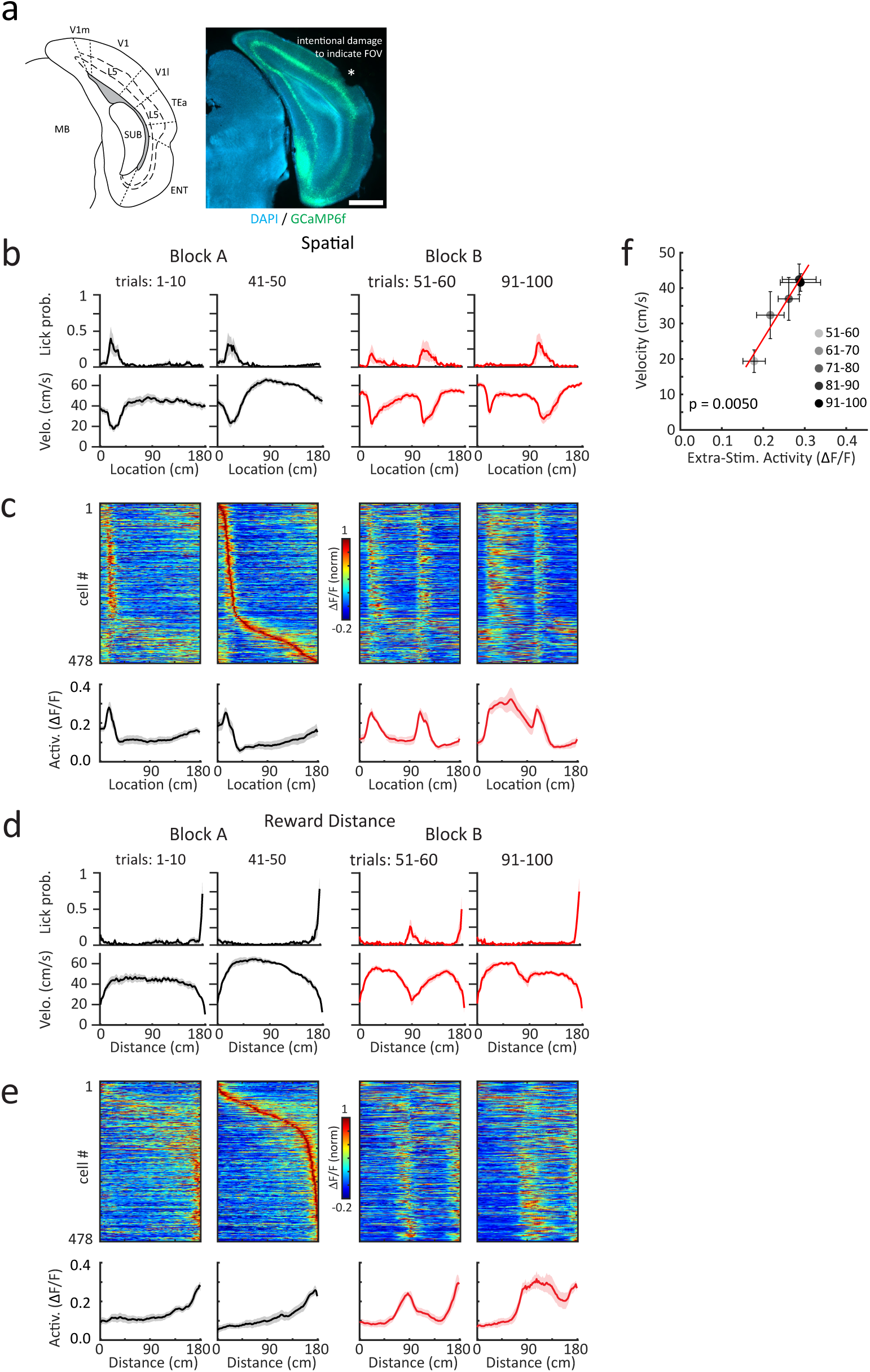
Histology, behavioral measurements and neuronal activities in HVA aligned to spatial position, and relationship between behavior and neuronal activities. **a,** Left, schematic illustration of cortical areas surrounding a typical imaging FOV. Right, histological image showing a representative FOV. The asterisk marks intentional tissue damage used to indicate the FOV location. Scale bar: 1 mm. **b,** Behavioral measurements (licking and running velocity) during the first and last 10 trials of each block, aligned to spatial location over the track (solid line, mean; shaded areas, mean ± SEM; n = 7 animals). **c,** Heatmap illustrating mean activities across all neurons during the behavioral phases shown in **b**, sorted by peak activity location during the last 10 trials of block A. Average neuronal activities (solid line, mean; shaded area, mean ± SEM; n = 7 animals) are shown below. **d, e**, Similar as in **b** and **c** but for behavior measurements and neuronal activities aligned to the distance from the last reward. **f,** Relationship between average velocity and average neuronal activity during the 1**–**2 s interval after S1 onset (trials 51–100, grouped into bins of 10 trials, indicated by color-coding; bars, ± SEM; red line, linear regression; *p* = 0.0050, n = 7 animals).

**Extended Data Fig. 2.**
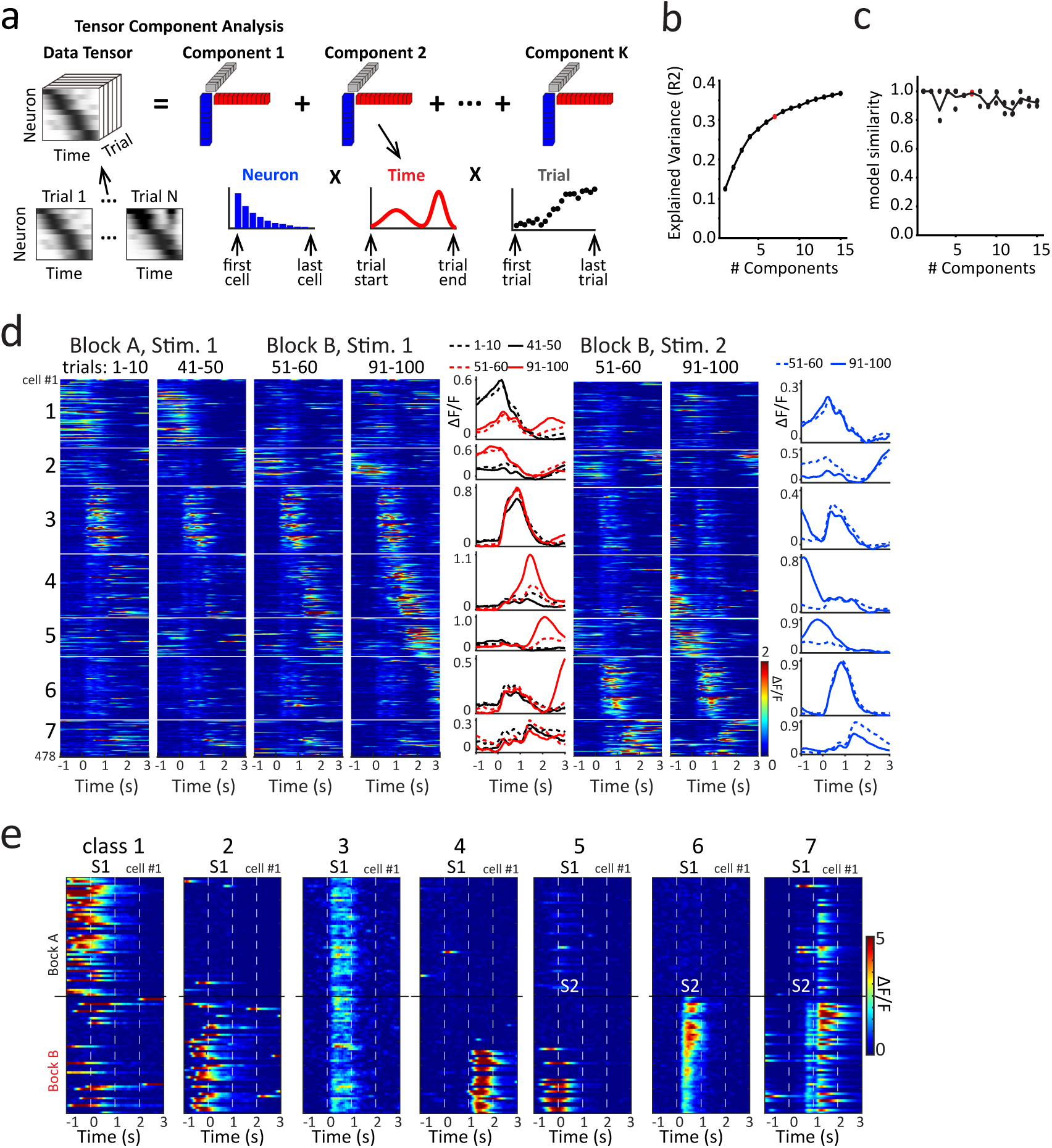
Tensor component analysis (TCA). **a,** Schematic illustration of tensor component analysis (TCA) as an unsupervised dimensionality-reduction method applied to large-scale neuronal recordings across trials. **b–c,** Fraction of explained variance of neuronal activity (**b**) and model similarity across different random initializations (**c**) as functions of the number of TCA components. The red dot indicates the component number (7) selected for analyses presented in Fig. 2. **d,** Mean activities of all neurons during the first and last 10 trials of each block, aligned to both stimuli and sorted according to cell’s component association from the TCA model shown in **a**. Average activities for neurons associated with each component are presented to the right. **e,** Activity of representative neurons associated with each TCA component across all trials in both blocks.

**Extended Data Fig. 3.**
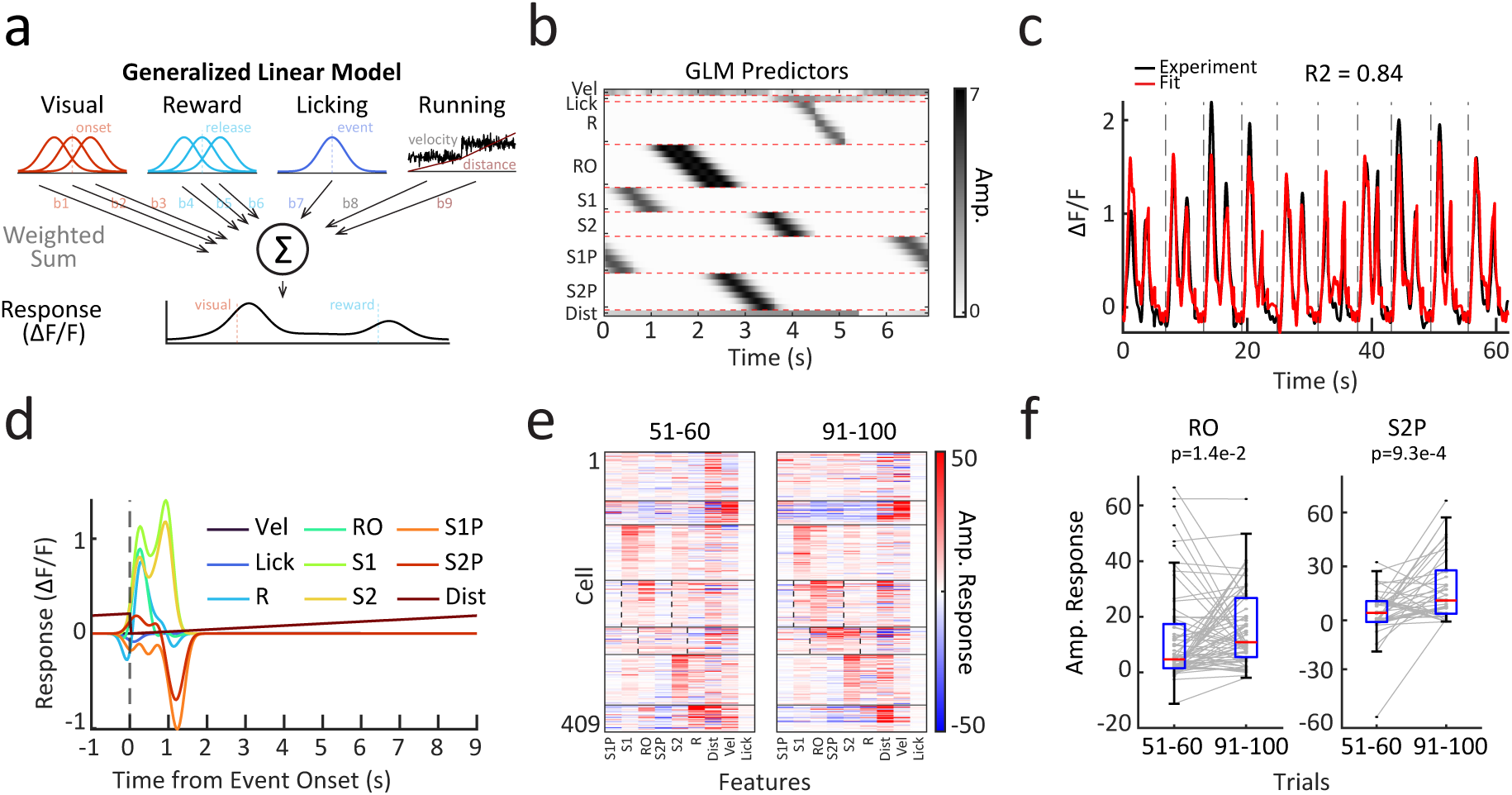
Generalized Linear Model (GLM). **a,** Schematic illustration of Generalized Linear Model (GLM), encoding behavioral and visual stimulus variables to predict neuronal activities. Discrete events (visual stimulus onset, reward release, licking) are smoothed using Gaussian kernels and then used as predictors in the linear model. **b,** GLM predictors for a representative trial, grouped by corresponding behavioral and visual stimulus variables (Vel: velocity; Lick: licking; R: reward; RO: reward omission; S1/S2: visual stimulus 1/2; S1P/S2P: prediction of visual stimulus 1/2; Dist: distance from last reward). **c,** Comparison of actual (black) and GLM-predicted (red) fluorescence (ΔF/F) traces for a representative neuron across 10 trials. **d,** GLM-fitted response profiles illustrating contributions of each behavioral and stimulus variable to predicted neuronal activity for the example neuron in **c**. **e,** GLM-fitted response amplitudes for individual behavior and stimulus variables across all neurons for the first 10 (left) and last 10 (right) trials of block B. Neurons are grouped by their TCA component associations and ordered consistently with Extended Data Fig. 2d. **f,** GLM-fitted response amplitude for variables “Reward Omission” and “S2 Prediction” compared between the first and last 10 trials of block B for neurons associated with the corresponding TCA components (one-tailed paired t-test, *p* = 0.014 and 9.3×10^−4^, respectively).

**Extended Data Fig. 4.**
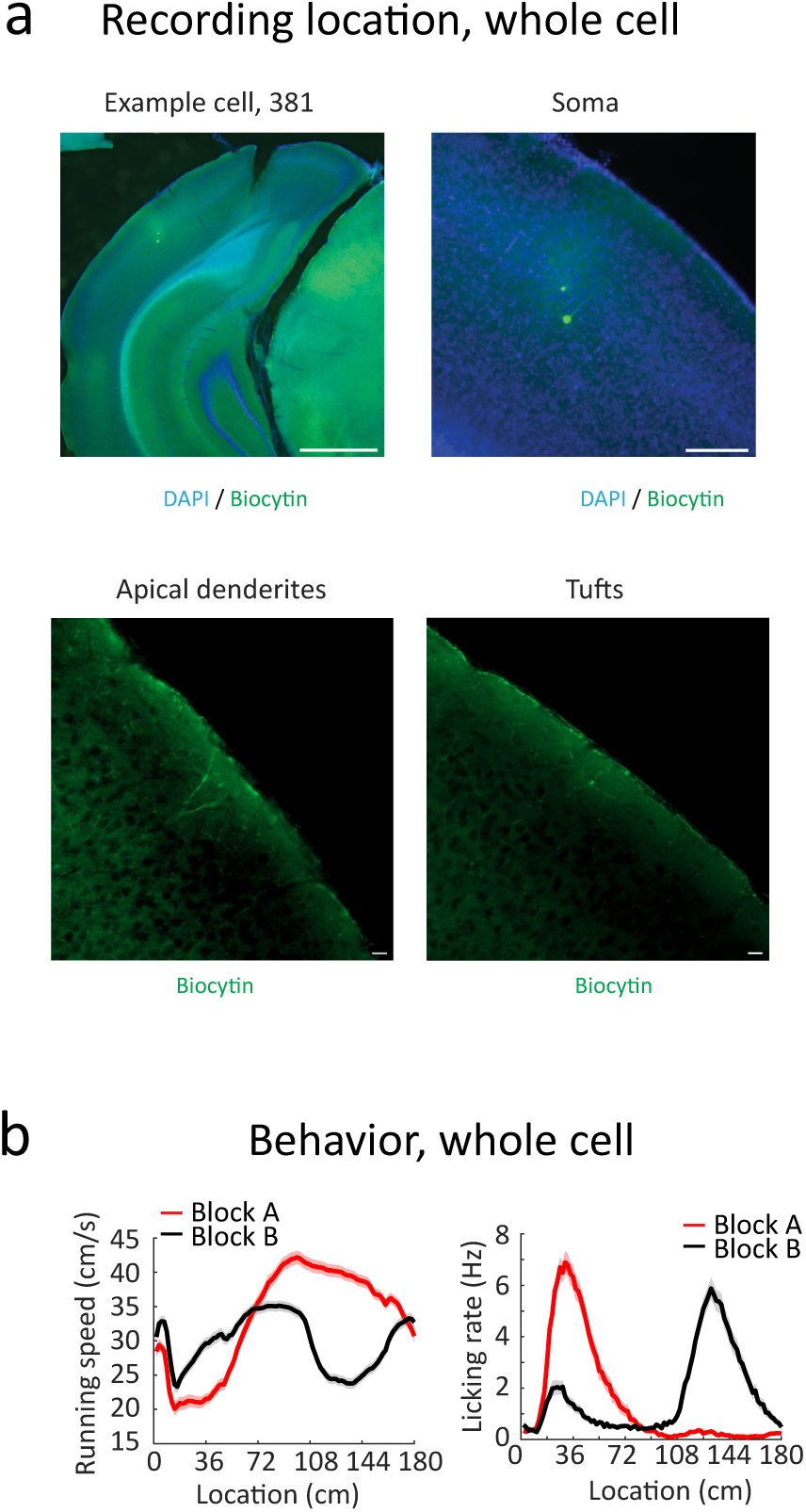
Recording locations and associated animal behavior. **a,** Representative histology confirming whole-cell recording locations. Top left: Low-magnification view of the recording site; Top right: expanded view of the soma; Bottom left: apical dendrites (100 μm posterior); Bottom right: tuft dendrites (200 μm posterior to the first panel). Note that dendrites extend posteriorly and L1 dendrites (right) are intact. Scale bars, 1000 μm (top left), 200 μm (top right), 20 μm (bottom left), 20 μm (bottom right). **b,** Behavioral metrics during whole-cell recordings. Animal speed (left) and licking (right) show pausing and increased licking at the newly rewarded stimulus (S2) location. Solid line, mean; shaded areas, mean ± SEM; n = 94.

**Extended Data Fig. 5.**
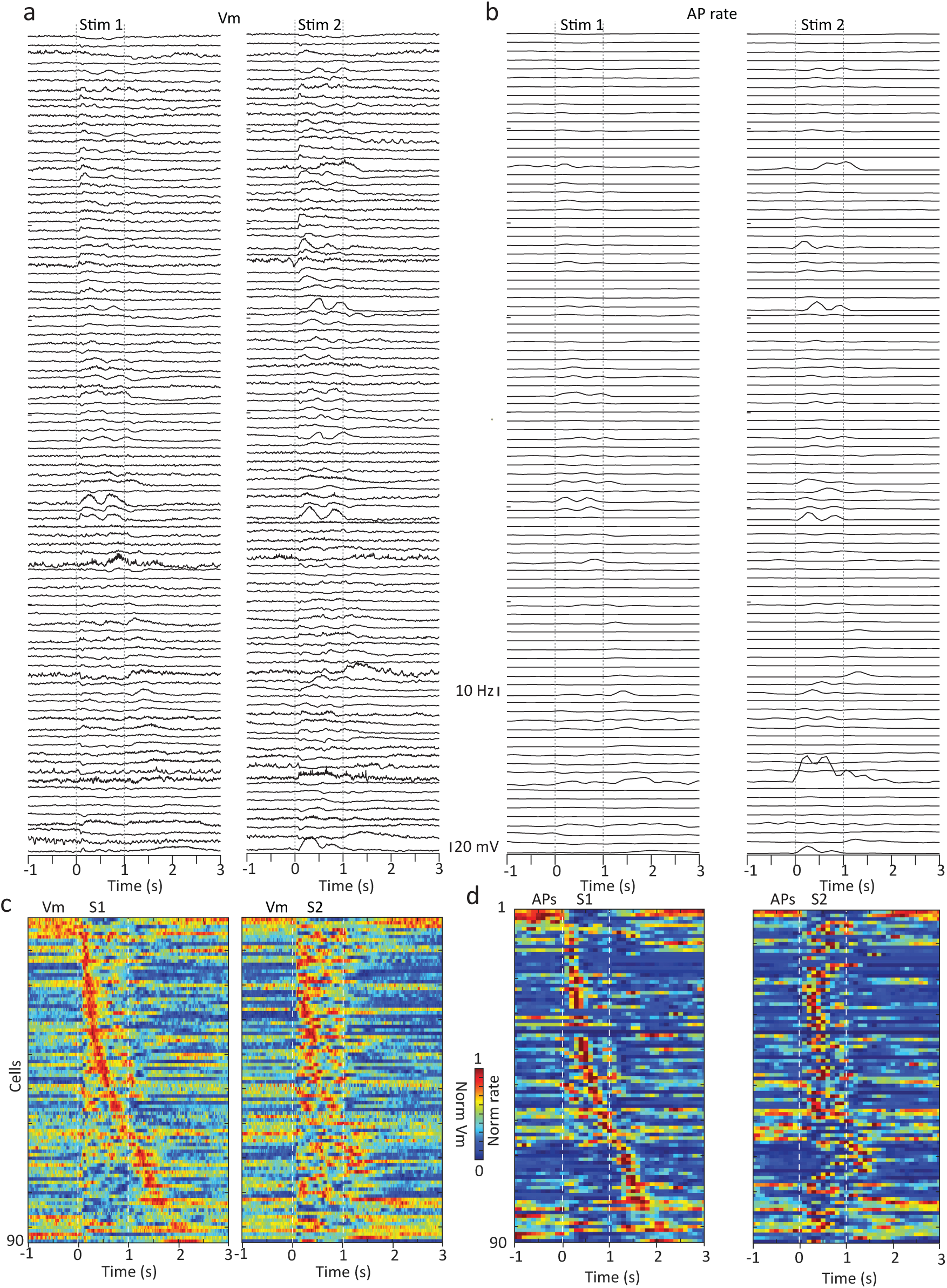
Membrane potential and firing rate responses to S1 and S2 in all recorded cells. **a**, Average Vm traces from all recorded cells in response to S1 (left) and S2 (right). **b**, Corresponding average AP firing rate traces for the cells shown in **a**. **c**, Heatmap representation of Vm responses from all cells to S1 (left) and S2 (right). **d**, Heatmap representation of AP firing rates for the corresponding cells in **c**. In all panels, cells are sorted by the peak response latency to S1 onset; the same sorting order is applied to S2 responses. All plots use data exclusively from block B before induction trials.

**Extended Data Fig. 6.**
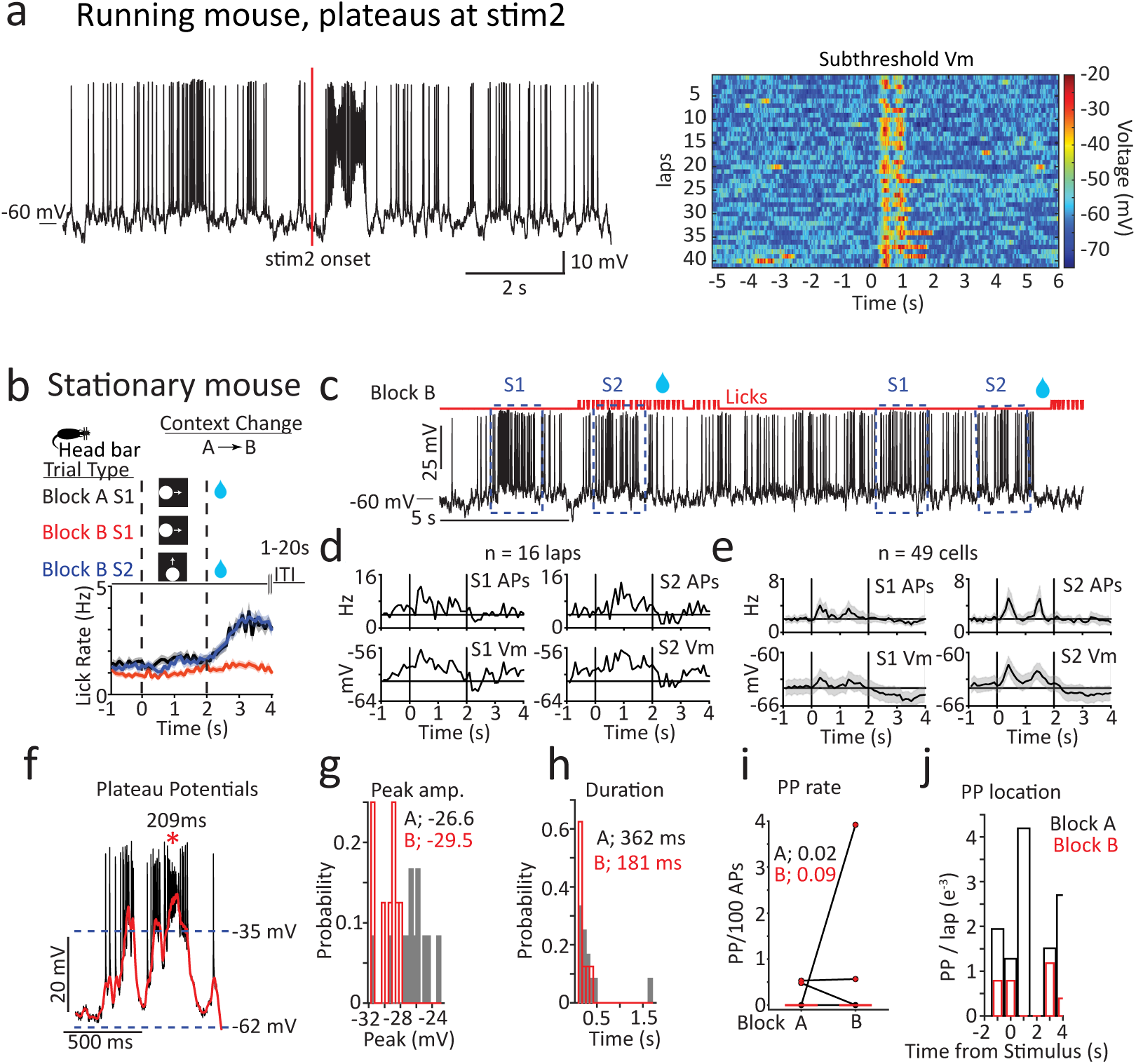
Spontaneous plateau in running and stationary mice. **a,** Example cell exhibiting consistent plateau potential firing following the contingency shift. Left, raw Vm trace showing two plateau potentials occurring after S2 onset. Right, heatmap of the same cell’s activity after the contingency shift, demonstrating consistent plateau potential generation aligned to S2 presentation in nearly all trials. **b,** Schematic of the stationary behavioral paradigm. Visual stimuli S1 and S2 (1 s each; two traversals per trial) were followed by a 1 s reward period and a 1 s vacuum period. In block A, S1 was paired with reward. In block B, reward was switched to S2, and S1 and S2 trials were presented in pseudorandom order with inter-trial intervals of 1–20 s. Population averages show licking behavior during the recordings for block A S1 (black), block B S1 (red), and block B S2 (blue), computed in 100 ms bins (n = 49 recordings from 28 animals). **c,** Representative membrane potential (Vm; black), licking events (red), and reward delivery (blue water symbol) from trials in block B. **d,** Average action potential (AP) rate and Vm for S1 and S2 from the neuron shown in **c** (n = 16 trials each). **e,** Population average AP rate and Vm for S1 and S2 (n = 49 neurons). **f,** Example spontaneous plateau potentials (PPs; raw Vm, black; smoothed Vm, red). Detection threshold denoted by blue dashed line. **g,** Distribution of PP amplitudes during block A (black) and block B (red). **h,** Distribution of PP durations during block A (black) and block B (red). **i,** PP rate normalized per 100 APs in block A and block B. **j,** PP rate versus time from visual stimulus onset in block A and block B (n = 20 PPs from 3 of 49 neurons with PPs).

**Extended Data Fig. 7.**
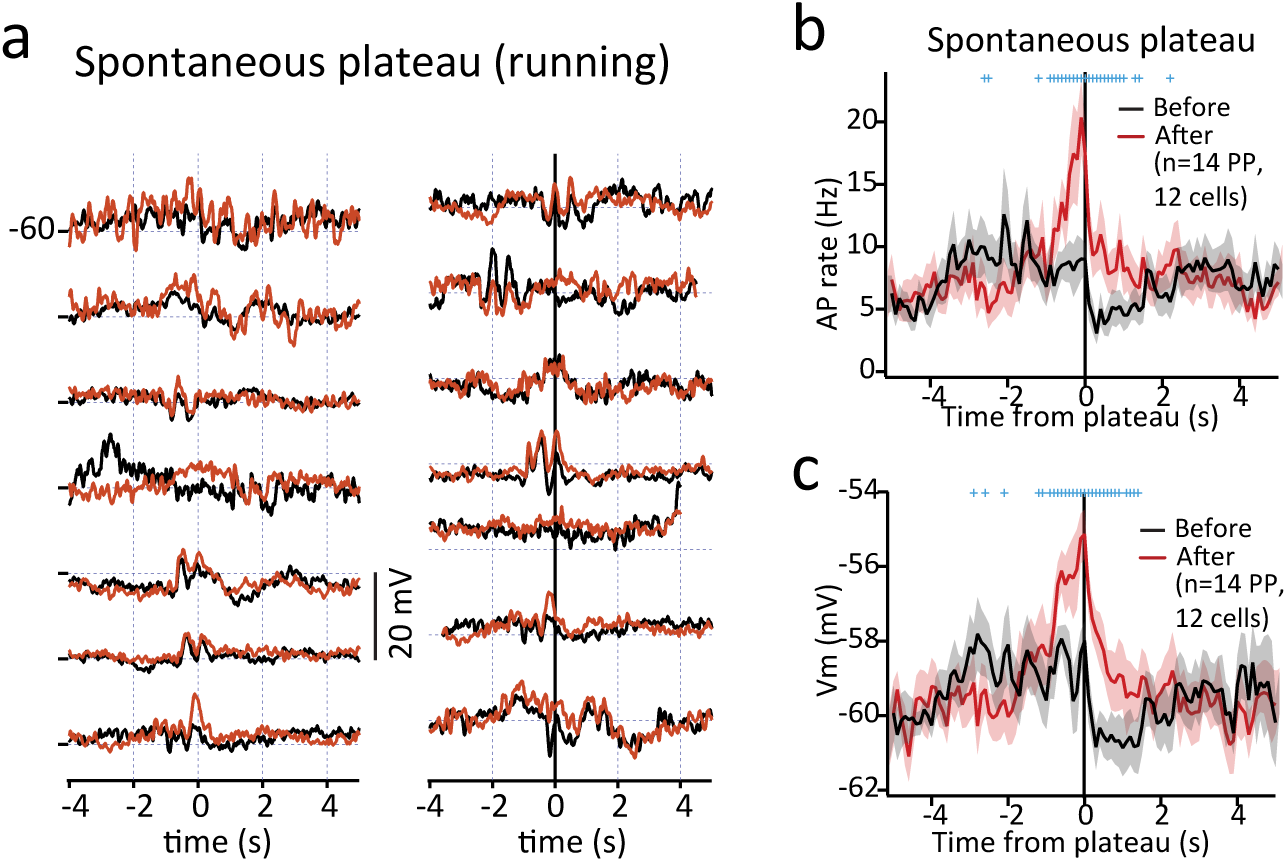
Analysis of spontaneous plateaus in running mice. **a,** Average Vm from all cells included in the spontaneous plateau analysis (black, before; red, after plateau). **b,** Average AP firing rate before and after the plateau potential. **c**, average Vm before (black) and after (after) the plateau potential. n = 14 from 12 cells. Solid line, mean; shaded areas, mean ± SEM; blue crosses indicate bins (100 ms) where *p* < 0.05 (two-way paired t-test).

**Extended Data Fig. 8.**
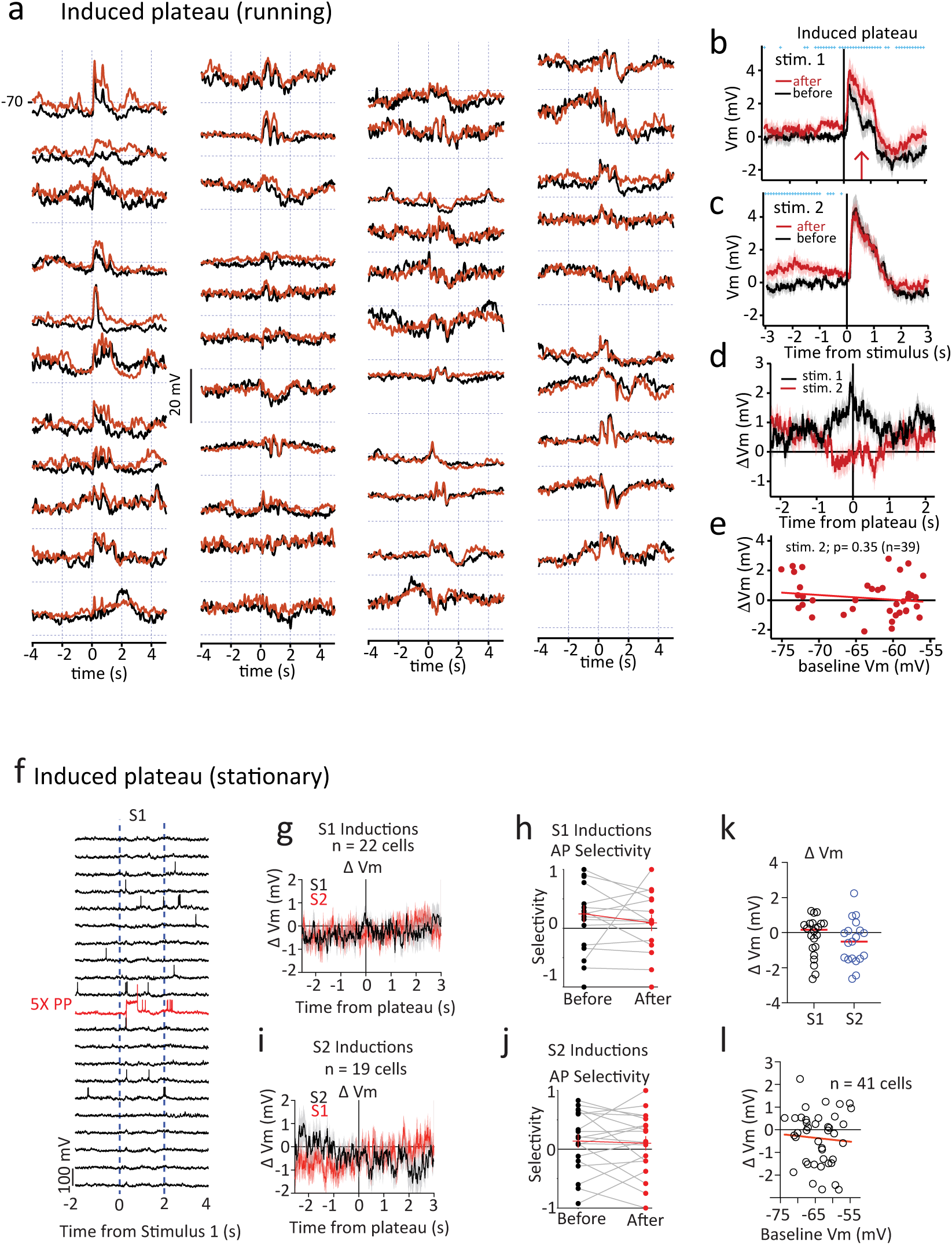
Analysis of induced plateaus in running and stationary mice. **a-e,** Data from running mice. **a,** Vm averages from all cells included in the induced plateau analysis for running mice. Traces aligned to S1 (black, before; red, after) **b,** Average Vm before (black) and after (red) the plateau potential around S1. **c,** Average Vm before (black) and after (red) the plateau potential around S2. **d,** the population average of the subtraction of before and after plateau induction traces; for period around S1 (black) or S2 (red). **e,** Correlation of the Vm change for S2 with the baseline Vm before induction. No significant correlation was found. For **b–d**, solid line, mean; shaded areas, mean ± SEM; blue crosses indicate bins (100 ms) where *p* < 0.05 (two-way paired t-test). **f-l,** Data from stationary mice. **f,** representative Vm traces from S1 trials, ten sequential S1 trials before, one of five S1 trials during PP induction, and ten sequential S1 trials after PP induction. **g,** Average change in Vm for S1 and S2 after PP induction to S1 (n = 22 inductions from 22 neurons; 10 ms bins; S1 black, S2 red). Mean ± SEM. **h,** AP selectivity index during the visual stimulus period (2–4 s) before and after PP induction for neurons with APs after induction (black, before; red, after). **i,** Average change in Vm for S1 and S2 after S2 PP induction (n = 19 inductions from 19 neurons; 10 ms bins; S2 black, S1 red). **j,** AP selectivity index during the visual stimulus period (2–4 s) before and after S2 PP induction for neurons with APs after induction (black = before; red = after). **k,** Average change in Vm during the visual stimulus period (2–4 s) for S1 after PP induction (n = 22 inductions from 22 neurons) and for S2 after PP induction (n = 19 inductions from 19 neurons). **l,** Change in Vm for S1 and S2 inductions versus baseline Vm for all neurons in stationary mice (n = 41 neurons).

**Extended Data Fig. 9.**
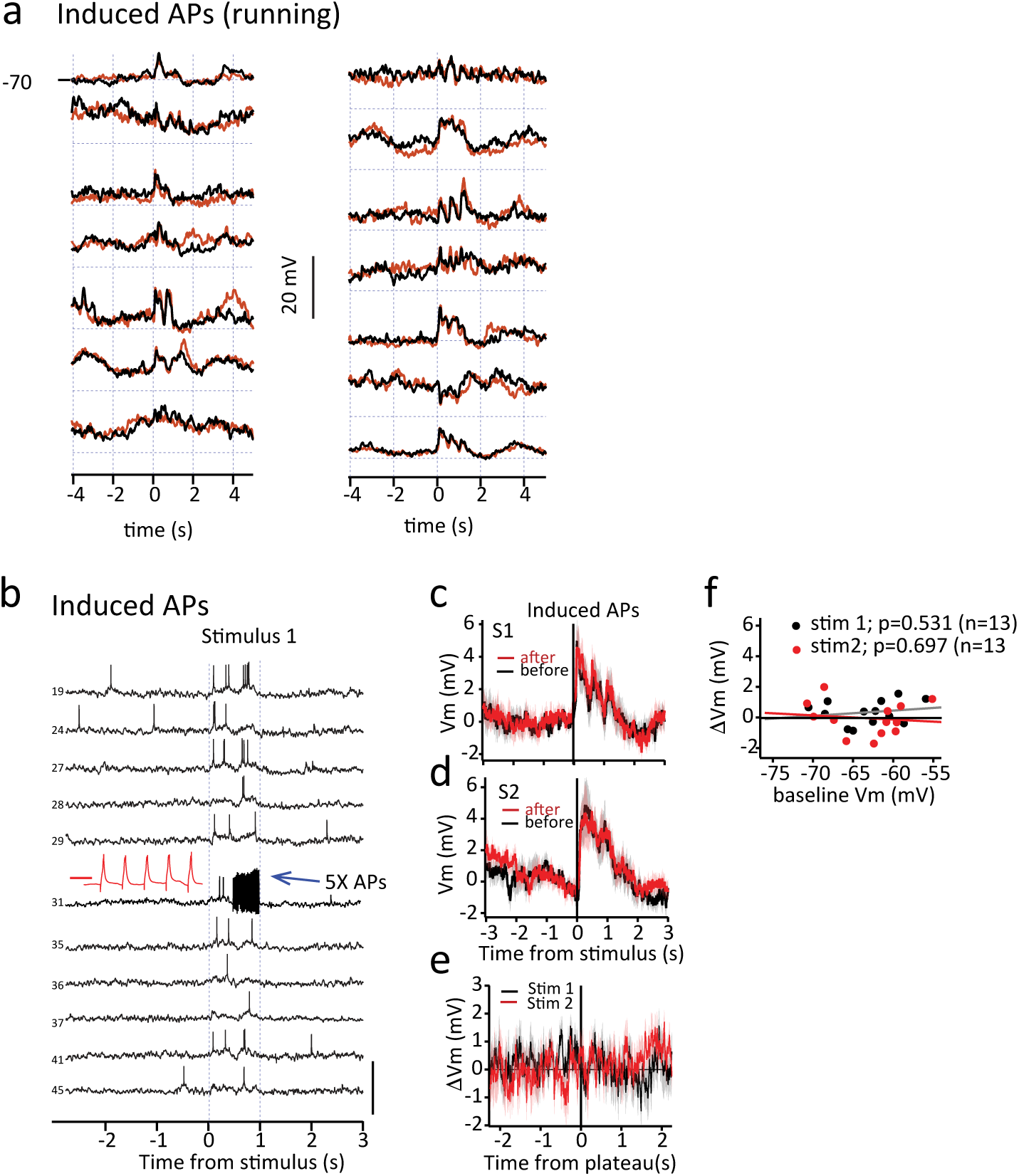
Analysis of action potential induction. **a-f,** Data from running mice. **a,** Traces from all cells included in the action potential induction analysis from running mice. **b,** Representative Vm traces for 5 selected laps before, during (1 of 5 laps shown), and 5 laps after induced APs. Inset: expanded view confirming AP generation by current injection. **c, d**, Average Vm responses aligned to S1 (**c**) and S2 (**d**) onset, for trials before (black) and after (red) AP induction. **e**, Average of subtracted Vm traces (after minus before AP induction) for S1 (black) and S2 (red). **f**, Correlation of Vm change with baseline Vm for S1 (black) and S2 (red). No significant correlation was found. In all relevant panels, blue crosses indicate bins (100 ms) where *p* < 0.05 (two-way paired t-test). *p*-values in **f** is from linear correlation.

**Extended Data Fig. 10.**
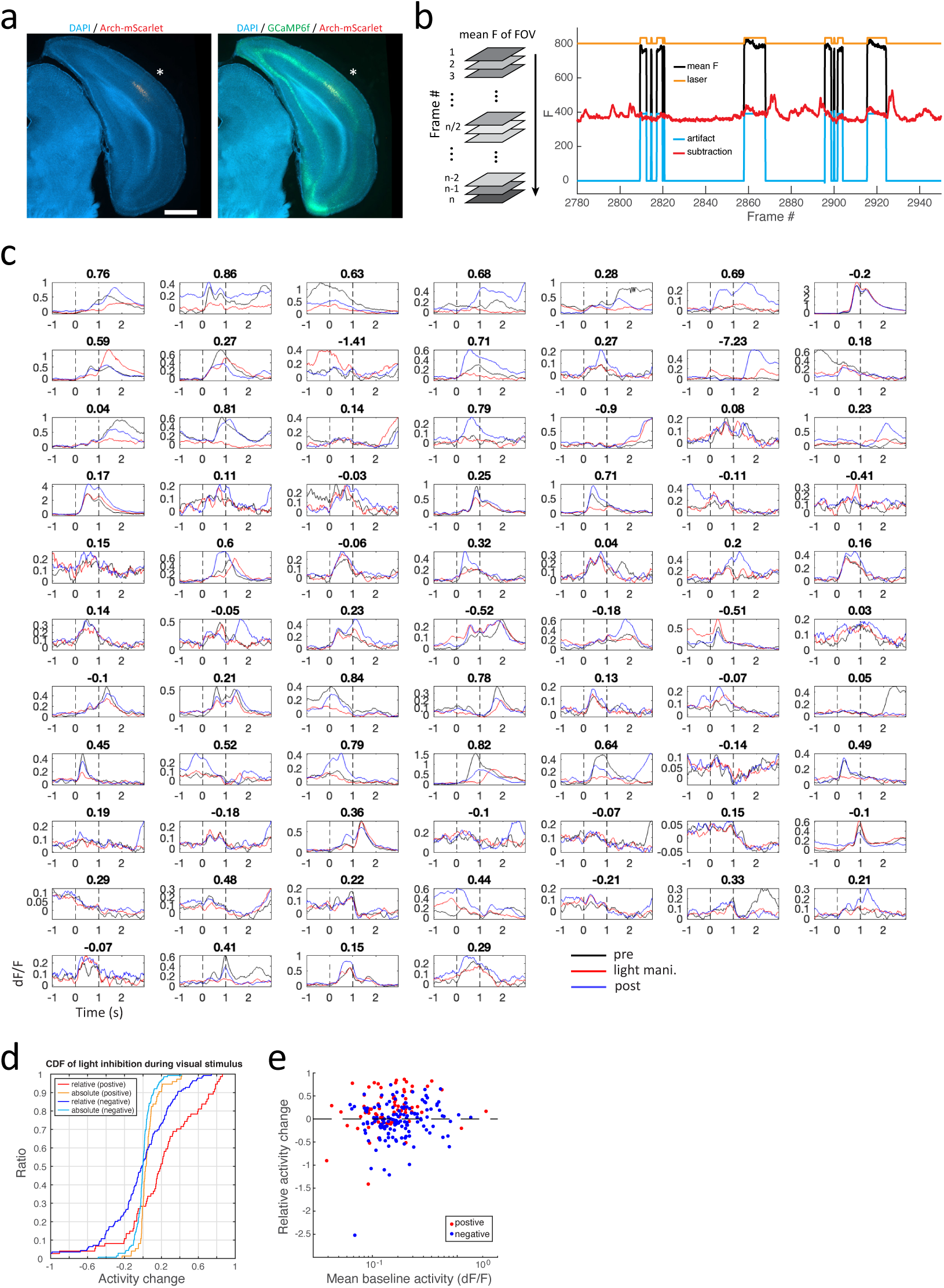
Effects of laser manipulation on neuronal activity. To assess the effects of laser manipulation, we performed an additional experiment on a separate day during block A, in which the laser was turned on for 1 s during presentation of S1 in the middle 30 trials, while the first and last 30 trials were without laser illumination in a subset of animals (n = 213 cells from 4 mice). **a**, Histology image showing the expression of Arch-mScarlet in the L5 of HVA. Asterisks indicate the FOV location. Scale bar: 1 mm. **b**, Illustration of laser artifact subtraction (Methods). Left, average fluorescence (F) across the entire field of view (FOV) for each individual frame. Right, example traces showing the mean F (black), laser trigger (orange), calculated artifact (blue), and the trace after subtraction (red). **c**, Averaged neuronal activity before (black), during (red), and after (blue) laser illumination aligned to S1 for all 74 cells with positive expression of Arch-mScarlet. Numbers indicate the relative activity change. **d**, Cumulative distributions of relative (dark color) and absolute ΔF/F (light color) changes in positive (red) and negative (blue, n = 139 cells) cells, respectively. Changes were calculated using activity measured during the light period. **e**, Scatter plot showing the relationship between relative activity change and mean baseline activity for individual positive (red) and negative (blue) cells, respectively.

**Extended Data Fig. 11.**
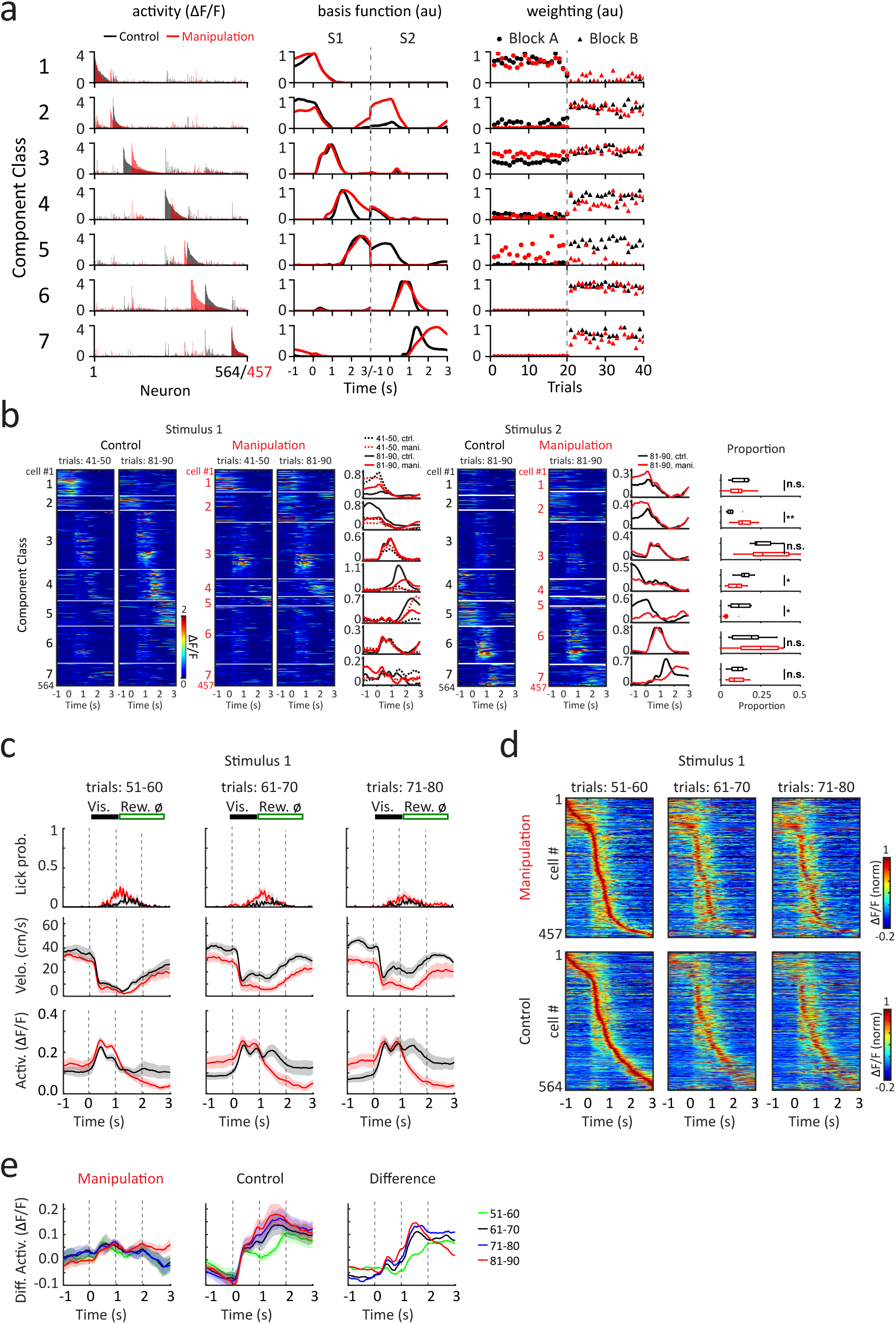
Behavior measurements and neuronal activities in HVA during optogenetic inhibition. **a,** Seven low-dimensional components identified by a rank-7 TCA model separately for control (black) and manipulation (red) group, analogous to Fig. 2a. Trial weighting factors are shape-coded according to the block assignment. **b**, Mean activity of all neurons from the manipulation and control groups, respectively, during the last 10 trials of block A and the first 10 trials of block B following optogenetic inhibition, aligned to both stimuli and sorted according to cell’s component association from the TCA model in **a**. Boxplots (right) illustrate the proportions of neurons most strongly associated with each TCA component (black, control group; red, manipulation group; n = 7 animals per group). **c,** Behavioral measurements (top, licking; middle, running velocity) and average neuronal activities (bottom; calibrated as in Extended Data Fig. 10b) during the first, middle and last 10 trials of block B under optogenetic inhibition, aligned to S1 (black, control group; red, manipulation group; solid line, mean; shaded areas, mean ± SEM; n = 7 animals per group). **d,** Heatmap illustrating mean activities across all neurons during the behavioral phases shown in **c**, sorted by peak activity time during the first 10 trials of block B under optogenetic inhibition (aligned to S1). **e,** Differences in average activity of the first, middle and last 10 trials of block B under optogenetic inhibition and the first 10 trials following inhibition, compared to the last 10 trials of block A (aligned to S1; left, manipulation; middle, control) and differences in average activity between control and manipulation groups during these behavioral phases (right).

## Methods

### Animals and surgery

All experimental methods were approved by the Institutional Animal Care and Use Committees of Baylor College of Medicine (Protocol 15–126). Wild type C57BL6 (WT; JAX #000664) male mice aged 8–14 weeks obtained from the Baylor College of Medicine Center for Comparative Medicine or Jackson Laboratory were used for all electrophysiology experiments. Rbp4-Cre (MMRRC: 031125-UCD) mice were crossed with Ai93 (JAX #024108) reporter mice in 10–18-week-old of either sex were used for two-photon imaging experiments. All surgical procedures were performed under ∼2% isoflurane anesthesia as previously described^44^. For imaging craniotomies, a 3.0-mm-diameter craniotomy was carefully created, centered at 3.6 mm mediolateral and − 0.1 mm anteroposterior relative to Lambda. A custom-made glass window was then implanted, consisting of three layers of glass (Potomac, #0): a top disk (outer/inner diameter, 4.5/2.5 mm), a middle ring(outer/inner diameter, 3.0/2.5 mm), and a 3.0 mm cover glass, glued together with UV-curable optical adhesive (Norland #71). The animals were singly housed after surgery and running wheels were added to their home cages. All mice were reared on a reversed 12/12 hr light/dark cycle with temperature (∼21 °C) and humidity (∼30-60%) controlled.For electrophysiology experiments, craniotomies for whole-cell and LFP recordings (∼0.5 mm in diameter) were centered at: 0.3 and 1.3 mm anterior, and 3.9 mm lateral to lambda, respectively. In a subset of animals, an optical fiber for optogenetic stimulation was implanted at 0.3 mm anterior and 2.0 mm lateral to lambda, angled 30° laterally to a depth of 2.0 mm. Following a minimum 7-day recovery period, animals were water-scheduled and habituated to head-fixation while running on a linear treadmill.

### Visual Stimulus

Visual stimuli were generated with PsychoPy v2022.2^45^in Python, and presentation was triggered by the animal’s position on a linear treadmill. All monitors were calibrated with a Calibrite Display Plus HL colorimeter. For whole-cell recordings a 15.6 inch ASUS MB16AHG LCD (1080p, 144 Hz) was positioned 15 cm from the left eye; a white disk (103 cd/m²) on a black background (0.08 cd/m²) subtended 65° in elevation (–28° to +37°) and 60° in azimuth (–27° to +33°), with a mean background luminance of 16.4 cd/m² under ambient room lighting. During motion trials the disk’s center moved horizontally at a constant elevation of +4.5°, sweeping 87° (–41° to +46° azimuth) at 174°/s; the upward motion spanned an elevation range from –33° to +41° at a speed of 148°/s, and the leftward and downward sweeps mirrored the rightward and upward motions, respectively, differing only in direction. For two-photon imaging an OLED ASUS MQ16AH (1080p, 60 Hz) was used, with the eye 12 cm from the screen center; a high contrast white disk (106 cd/m²) on a 0.03 cd/m² background (in a darkened room) matched the angular size of the whole-cell disk. Each motion trial consisted of two consecutive 0.5 s sweeps (total 1 s); in the imaging paradigm a rightward moving disk was presented at the 10 cm treadmill mark, and an upward moving disk at the 100 cm mark.

### Behavioral Apparatus and Training

Behavioral control and data acquisition were managed by a MATLAB GUI interfacing with a Bpod Finite State Machine (r1.0, Sanworks). Mice were trained on a linear treadmill (∼180 cm long) consisting of an empty velvet fabric belt. Animal running velocity was measured by a rotary encoder attached to a treadmill wheel axle, and distance was calculated by integrating velocity within a lap. Photoelectric sensor at the track’s beginning detected and reset the lap location. A 10% sucrose solution reward (∼3.3 μL per drop) was delivered via a custom-made lick port controlled by a solenoid valve (Lee Valve, LHQA1231220H_B), and licking was detected by an optical sensor (Panasonic, FX-301H). This Bpod system delivered position-dependent visual stimuli and rewards. A separate custom Arduino-based (Teensy 3.5) module controlled position-dependent intracellular current injection. All behavioral data were digitized at 20 kHz by a PCIe-6343, X-series DAQ system (National Instruments) using WaveSurfer software (wavesurfer.janelia.org).

Following a minimum one-week recovery period, mice underwent water scheduling, maintaining their body weight above 80% of their pre-restriction weight. Water was provided during experimental sessions and supplemented daily. Animals were handled by the experimenter for approximately 30 minutes daily for five days. Treadmill training began with 20-minute sessions where rewards were delivered every 30 cm to encourage running. The reward interval was incrementally increased by 30 cm every 10 trials until it reached 180 cm (one lap), at which point the reward was fixed at that location. By the second training day, the reward location was typically fixed, and sessions were extended to 40 min. From the third day onward, sessions were limited to 1 h. Once animals reliably completed 50 laps per session, familiar visual stimuli (S1) were introduced at 10 cm, paired with the reward. Mice were considered ready for task switching when they consistently completed 150 laps within 1 h for three consecutive days. On the subsequent day, novel visual stimuli (S2) were introduced at 100 cm starting from the 51st lap, and the reward contingency was shifted from S1 to S2. For the electrophysiological recordings, Following the successful establishment of a whole-cell patch, a stable baseline recording was acquired for approximately 5 minutes. Immediately after this baseline period, the reward contingency was shifted.

### Two-photon imaging

Two-photon imaging was performed using a custom-built microscope (Janelia MIMMS2.0). GCaMP6f in cortical L5 neurons, typically at a depth of 450–550 µm below the pia, was excited at 920 nm using a Ti:Sapphire laser (Chameleon Ultra II, Coherent) with a typically power of 50–100 mW measured under the objective (×16, 0.8 NA, Nikon). Emitted florescence was separated by primary (FFF705-Di01) and secondary (FF573-Di01) dichroic mirrors (Semrock) and further filtered by a FF01-515/30 (Semrock), a BG39 (Schott) and an additional FF01-515/30 bandpass filter before detection with a GaAsP photomultiplier tube (11706P-40SEL, Hamamatsu). For the red detection arm, a FF02-617/73 bandpass filter (Semrock) was used. Images (512×512 pixels) were acquired at ∼30 Hz using ScanImage software (Vidrio Technologies). Imaging fields of view (∼300×300 µm) were selected within the lateral and posterior quadrant of the cranial window.

### Optogenetic manipulation

*AAV-hSyn1-Flex-ArchT-IRES-mScarlet* (control: *AAV-hSyn1-Flex-tdTomato* or saline) with a titer of 7.4×10^12^ was injected into the L5 (700 and 500 µm below the pia) of the higher visual area (HVA; anteroposterior: ∼0.2 mm to Lambda, mediolateral: ∼3.9 mm from the midline) in *Rbp4-cre:: Ai93* transgenic mice prior to glass window implantation. For each injection depth, 10 pulses (2.5 nl per pulse) were delivered at 30 s intervals, with a 5 min between depths and an additional 10 min pause before complete withdrawal of the pipette. Approximately 3 weeks after injection, mice were placed under water-restriction and behavioral training commenced. Data collection was performed roughly one-month post-injection.

For simultaneous imaging and optogenetic inhibition, the primary dichroic mirror was replaced with a Di02-R561 (Semrock). ArchT was activated using a 594 nm laser (Cobolt), the beam of which was expanded to uniformly fill the back aperture of the objective. Laser power was set to 0.4 mW, measured after the objective, and controlled by a custom-built Arduino-based module (Teensy 3.5). During light delivery trials, the laser shutter opened immediately following the S1 offset and was subsequently modulated based on the mouse’s running speed (close when velocity <5 cm/s for periods exceeding 1 s). The laser shutter was closed at the onset of S2.

To correct fluorescence artifacts caused by mScarlet excitation under 594 nm laser illumination, we first computed the mean fluorescence (F) across the entire imaging FOV for each frame (Extended Data Fig. 10b). Periods of laser illumination were identified using the laser trigger signal. For each illumination period, artifact values for the first and last frames were determined as the differences between those frames and the frame immediately before or after, respectively. Artifact values for the intermediate frames were assigned a constant value, calculated as the mean difference between the second and penultimate illuminated frames and their respective corresponding neighboring frames (two frames before or after), averaged across all illumination periods. Finally, the calculated artifact was subtracted from the raw F of each ROI for each imaging session.

### In vivo electrophysiology

We performed in vivo whole-cell patch-clamp recordings from neurons in the deep layers (550–750 µm) of higher visual areas. A patch pipette (8–12 MΩ) was angled at 22.5° from vertical and advanced using a micromanipulator (Luigs & Neumann, SM-10). Neuronal depth was determined from the micromanipulator’s travel distance relative to the pial surface, which was identified electrically upon contact. For simultaneous local field potential (LFP) recordings, a second electrode (2–3 MΩ) was positioned at a 45° angle. Whole-cell recordings were performed using elongated-taper borosilicate pipettes (8–12 MΩ) filled with an internal solution containing (in mM): 134 K-gluconate, 6 KCl, 10 HEPES, 4 NaCl, 0.3 Mg-GTP, 4 Mg-ATP, and 14 Tris-phosphocreatine. For subsequent morphological reconstruction, 0.2% biocytin was included in a subset of recordings. To approach neurons, positive pressure (∼10 psi) was applied to the pipette to penetrate the dura, after which it was reduced to ∼0.25 psi. Cell contact was identified by a characteristic and reproducible increase in tip resistance. All recordings were made in current-clamp mode with a Dagan BVC-700A amplifier and digitized at 20 kHz using a National Instruments DAQ (PCIe-6343) and WaveSurfer software. Series resistance was compensated using the amplifier’s bridge balance circuit, and recordings were discarded if it exceeded 70 MΩ.

### Stationary Whole-Cell Experimental Procedures

Animal surgery was similar to described above. Mice were trained to receive a 10% sucrose reward (3.5 μL) following presentation of two consecutive 1-s sweeps (total duration 2 s) of a rightward-moving white disk on a black background (S1) while head-fixed on an air table. The reward was available for 1 s after stimulus presentation and was subsequently removed by vacuum. The interstimulus interval was randomized between 1 and 20 s. Training continued for 5–10 days until animals consistently consumed ≥70% of rewards during 60-min sessions (150–200 trials). Supplemental water was provided as needed to maintain stable body weight.

Visual stimuli were generated using RPG, a Python package for the Raspberry Pi ^46^,and presented on a 27-inch LG MP89HM-S IPS LCD monitor (1080p, 60 Hz) positioned 15 cm from the left eye. The moving disk traversed equal distances across the monitor for S1 (leftward) and S2 (upward) conditions, covering ∼44° of the visual field and elevated ∼20°. The reward valve, lick sensors, and stimulus triggers were controlled using a Bpod finite-state machine (r1.0, Sanworks).

Following training, a craniotomy was performed, and in vivo whole-cell patch-clamp recordings were obtained the next day. Recordings targeted deep layers (550–750 μm below the pial surface) of higher visual areas. Elongated-taper borosilicate glass pipettes (tip resistance: 9.52±0.15 MΩ, SEM) were positioned at a 30° angle from vertical using a Luigs and Neumann micromanipulator. Whole-cell recordings were conducted under similar conditions as running experiments. Each recording included an initial baseline period (block A, ∼25 trials), followed by a second period (block B, ∼25 trials) in which S2 was introduced and the rewarded stimulus was switched from S1 to S2. Trial types were pseudorandomized, with no more than three consecutive presentations of the same type. For induction trials, S1 and S2 stimuli were presented in sequence with a fixed intertrial interval, and either S1 or S2 was paired with a 500-ms depolarizing current injection (∼1–1.2 nA) delivered 300 ms after stimulus onset. The average recording duration was 30.2±1.7 min (SEM), comprising 140±8.2 trials (SEM) across 49 neurons recorded in the stationary condition.

### Cell inclusion criteria for in-vivo electrophysiology

The final dataset for plateau analysis included only cells with recording durations sufficient to complete the entire induction protocol (∼10 trials after the induction) and was restricted to putative pyramidal neurons. Putative interneurons were identified and removed from analysis using an unsupervised k-means clustering protocol. The clustering was based on five key electrophysiological parameters: Afterhyperpolarization (AHP): The mean AHP amplitude from 500 spikes (or the maximum available) within the recording segment from the 2-min mark onward. Baseline Membrane Potential (Vm): The average Vm in the 500 ms window (0.75 s to 1.25 s) preceding the first visual stimulus. Overall Firing Rate: The mean firing rate across the entire recording. Plateau Firing Rate: The mean plateau rate during identified plateau potentials. Visual Stimulus Response: The average Vm during the 1-sec period of the first visual stimulus. Prior to clustering, these features were standardized (z-scored) and weighted to emphasize the discriminative power of AHP (weights: 1.0, 0.3, 0.3, 0.1, and 0.1, respectively). The optimal number of clusters was determined to be four (k = 4) using the elbow method. A distinct cluster (20% of cells) with a positive AHP centroid, a characteristic of interneurons, was identified.

Consequently, all cells within this cluster were excluded from the final BTSP analysis. AHP amplitudes in individual neurons were quantified as the difference in the minimum (2 ms average) Vm amplitude in the period preceding the average AP (100 ms before) and the minimum (2 ms average) Vm after the AP (50 ms after) (AHP amplitude: interneuron group, −1.5 pm 0.2 mV; n = 17; pyramidal neurons +1.1 pm 0.2; n = 43).

### Data analysis for in-vivo electrophysiology

As described previously^7,47^, we performed a trial-by-trial Vm correction by identifying the action potential (AP) threshold, calculating its offset from −50 mV, and subtracting this offset from the entire trace. For plateau potential analysis, APs were first excised by removing data 1.5 ms before and 3.5 ms after the threshold; the resulting gap was filled using linear interpolation. This AP-free trace was then smoothed with a moving average (20.1 ms window), and plateaus were defined as events in the smoothed trace crossing a −35 mV threshold. We characterized each plateau by its peak (maximum Vm of the AP-free trace), duration (time above −35 mV), and location (average animal location during the event). The plateau rate was calculated as the total number of plateaus divided by the total AP count within block A or block B.

For the analysis of spontaneous plateaus, induced plateaus, and induced APs, we first applied the Vm correction and AP removal (excision and interpolation) procedures described above. We then averaged “before” and “after” induction trials, selecting an equal number of trials (typically 10) from all available trials within the same behavioral block based on the smaller available count.

The average “before” trace was subtracted from the average “after” trace to generate a difference trace, isolating the induction effect. The change in membrane potential (ΔVm) was quantified as the average of this difference trace during the visual stimulus period or a 1 second period around the center of a spontaneous plateau. Baseline Vm was calculated as the average Vm for the 1 second period before S1. In the average traces shown in Extended data figure 8b,c and 9c,d the baseline value has been subtracted from the individual traces before averaging. The selectivity index was calculated from AP firing rates as (S1−S2)/(S1+S2).

### Data analysis for Two-photon imaging

Acquired two-photon images were motion-corrected using Suite2P^48^(Python version, http://github.com/MouseLand/suite2p). Regions of interest (ROIs) were determined and time-series fluorescence data extracted by automatically detection in Suite2P, followed by manual curation. Neuropil subtraction was disabled. Further analyses were conducted using custom MATLAB code. Raw fluorescence signals underwent baseline correction using a 30-s moving window, after removing signals exceeding 1 s.d. above baseline. The signals were then converted to ΔF/F ((F−F0)/F0), where F0 was defined as the median of a 30-s moving window of baseline. And ΔF/F traces were further smoothed with 0.2-s moving window.

### Tensor Component Analysis (TCA)

We performed tensor component analysis (TCA) as an unsupervised dimensionality-reduction method to decompose large-scale neural recordings into low-dimensional components representing coherent neural dynamics. TCA identifies neural components described by three sets of factors: neuron factors capturing groups of co-activated neurons, temporal factors capturing the within-trial activity patterns, and trial factors capturing across-trial dynamics (reflecting gradual changes such as learning) ^49^ . We use the Python package “tensortools” with the “nonnegative canonical polyadic decomposition by hierarchical alternating least squares” (“ncp_hals”) algorithm, which constrains all factor entries to be non-negative.

We constructed a single three-dimensional data tensor with dimensions neurons × trial time × trials encompassing all recorded neurons from 7 mice (n = 478 cells) and all trials from two task blocks (block A and block B). Each trial was represented by an 8-sec time series capturing responses aligned around two stimuli. Specifically, we concatenated the activity segments aligned to S1 (from 1 s before to 3 s after stimulus onset) and S2 (also from 1 s before to 3 s after onset), yielding a unified activity vector for each trial. For trials in block A (which had only S1 and no S2), the second 4-sec segment corresponding to S2 was padded with zeros to maintain a consistent length of 8 seconds. This neuron-by-time-by-trial tensor was the input to TCA.

We fit TCA models across a range of ranks (numbers of components) from 1 to 15, with 4 independent random initializations per rank to form an ensemble of fits. To determine the optimal rank, we evaluated two metrics: (1) the fraction of variance explained (analogous to an R² metric in linear regression), quantifying how well the low-dimensional factors reconstructed the original neuronal data, and (2) the model similarity across different random initializations. As the number of components increased, the fraction of variance explained initially grew but plateaued after six components (Extended Data Fig. 2b). Within this range, the decomposition exhibited high stability at rank 7: the similarity of components from different random initializations exceeded ∼0.95 (Extended Data Fig. 2c). This high consistency indicated that seven components could be reliably identified regardless of initialization. Based on these metrics, we selected a rank-7 TCA model for all subsequent analysis.

Following extraction of the rank-7 TCA model, we rescaled the factors for clearer interpretation. There is an inherent scaling invariance in TCA decompositions (scaling one factor can be exactly compensated for via the inverse scaling of another), so we imposed a standardized scaling approach. We normalized each component’s temporal factor and trial factor to the range [0, 1] by dividing each factor’s maximum value, and correspondingly multiplied the neuron factor by that same value to preserve the decomposition. This normalization placed the temporal profiles of all components on a comparable scale and allowed the neuron-factor weights to reflect the true amplitude of each component’s contribution to neural activity. After rescaling, the magnitude of a given neuron’s weight in a component’s neuron factor can be interpreted as the contribution of that component to the given neuron’s activity pattern. Finally, we assigned each neuron to the component in which it had the highest neuron-factor weight, effectively clustering the 478 cells into 7 groups. Each resulting group thus represents a cell assembly characterized by similar within-trial activity patterns and across-trial dynamics, as defined by the corresponding TCA component.

### Generalized linear model analysis

We applied a generalized linear model (GLM) to quantify the tuning of each neuron’s calcium activity to behavioral and stimulus variables, and to examine how the tuning changed between the early and late phases during learning (block B). We modeled the calcium fluorescence signal (ΔF/F) of each neuron independently using a GLM with a Gaussian error distribution (identity link function). This cell-by-cell GLM approach provides a set of specific coefficients for each neuron, capturing each neuron’s unique tuning profile. Unlike tensor component analysis (TCA), which identifies population-level covariance patterns, the GLM explicitly separated the contributions of individual behavioral and stimulus variables to each neuron’s activity.

Behavioral and stimulus predictors were constructed at the imaging frame rate (30 Hz) (Extended Data Fig. 3). Licking events were downsampled to 30 Hz and represented as a binary time series (lick present/absent per frame), then convolved with a Gaussian kernel (σ = 5 frames) to produce a smooth licking regressor. Reward delivery events were similarly downsampled and convolved with a series of 7 Gaussian kernels (σ = 5 frames), each temporally offset by increments of 5 frames, to span the 1-second window (30 frames) following reward release. For unrewarded stimulus presentation (reward omission), we defined a binary event at the expected reward time (1 s after the unrewarded stimulus onset) and convolved it with the same set of 7 Gaussian kernels, capturing neural responses to omitted rewards. Visual stimulus for each of the two stimulus was encoded as a 0.5 s long event (15 frames from stimulus onset) and convolved with 4 Gaussian kernels (σ = 5 frames) evenly spaced across 0.5 s (15 frames) until the offset of stimulus, to capture the transient visual-evoked responses. To capture predicative activity prior to stimulus, we also included stimulus prediction predictors: for each stimulus, the 0.5 s stimulus onset event was convolved with six Gaussian kernels (σ = 5, each offset by decrements of 5 frames) covering the 1 s interval before stimulus onset. Continuous variables including the animal’s running velocity and the distance traveled since the last reward are both sampled at 30 Hz and aligned to the imaging frames. For numerical stability and to allow comparison of coefficient magnitudes, all predictor variables were normalized to the range [0, 1] before model fitting.

GLMs were fitted using the MATLAB implementation of the “glmnet” package. Separate GLMs were fitted for each neuron and for each phase of Block B, using the first 10 trials and the last 10 trials. We used an elastic net regularization with an equal mix of L1 and L2 regularization (α = 0.5). The regularization strength for each model was selected via five-fold cross-validation, choosing the value that minimized the cross-validated error. Model performance was quantified by the explained variance (R²) of the GLM predictions (Extended Data Fig. 3c). For subsequent analyses, we conservatively included only neurons with an R²>0.1 (i.e. >10% variance explained) in both first 10 and last 10 trials.

To quantify each neuron’s selectivity for specific variables, we used the fitted GLM coefficients to simulate the neuron’s response during a standardized “virtual lap”. In this virtual lap, the behavioral trajectory was fixed to the average profile (e.g. mean running speed and licking rate) and visual stimuli and rewards were positioned at the same spatial locations as in actual experiment. Using the same encoding and normalization as for real data, we generated the full set of predictor variables for this virtual lap. We then computed the model’s predicted calcium trace when including predictors associated with only a single variable at a time (setting all others to zero). The area under the curve (AUC) of each variable-specific predicted response (Calcium trace) was taken as a measure of the neuron’s response magnitude driven by that variable. This AUC therefore quantifies the cell’s selectivity for each behavioral or stimulus variable during the given phase (Extended Data Fig. 3d).

## Acknowledgements

We thank R. Chitwood for technical assistance. We thank members of Magee lab for useful discussions. This work was supported by the Howard Hughes Medical Institute and the Cullen Foundation.

## Contributions

K.X., Y.L., B.S. and J.C.M. designed the research. K.X. performed electrophysiological recordings in running setup. Y.L. performed imaging recordings and optogenetic manipulation. B.S. performed electrophysiological recordings in standing setup. G.L. designed and implemented the computational model. K.X., Y.L., B.S., G.L. and J.C.M. analyzed the experimental data. K.X., Y.L., G.L. and J.C.M. wrote the manuscript.

